# Study of the immunogenicity, efficacy and safety of recombinant RBD SARS-CoV-2 vaccine with CpG adjuvant in rodent, non-rodent and Maccaca fascicularis using Indonesian Strain Virus

**DOI:** 10.1101/2024.12.18.629000

**Authors:** Nur Amalia Limeilati, Reviany V. Nidom, Trilokita Tunjung Sari, Nurvita P. Kusumarahayu, Siti V. Fransiska, Setyarina Indrasari, Afrillia N. Garmana, Dhyan K. Ayuningtyas, Gembong S. Nugroho, Darsono, Maharani, Acep R. Wijayadikusumah, Elin Yulinah, I Ketut Adyana, Astria N. Nidom, Neni Nurainy, Chairul A. Nidom

**Author notes:** Equally (Neni Nurainy, Chairul A. Nidom).

## Abstract

SARS-CoV-2 is the leading cause of the COVID-19 pandemic that causes acute respiratory syndrome, emerged in late 2019, and was declared a global pandemic on March 11^th^, 2020. A safe and effective vaccine that prevents SARS-CoV-2 infection or minimize SARS-CoV-2 disease burden is needed. However, in 2021, Low- and middle-income countries (LMICs) face challenges regarding supply of COVID-19 vaccines. Indonesia, as a public sector vaccine manufacturing in developing countries was developed COVID-19 vaccine using a platform based on recombinant subunit proteins, a Receptor Binding Domain (RBD) of SARS-CoV-2 formulated with combination of Alhydrogel and CpG Oligodeoxynucleotides 1018 (CpG). In this study, we report the preclinical study including immunogenicity, toxicity and efficacy of vaccine in animal models. The vaccine immunogenicity tested in mice and non-human primates, the toxicity was done in rodents and non-rodents and challenged study and efficacy was done in non-human primates (NHPs) model. The animal model was vaccinated intramuscularly (IM). The serology result in mice and non-human primates showed significant antibody titers and neutralizing antibody responses compared to the RBD formulations adjuvanted with Alhydrogel only. Safety study in Wistar rats and New Zealand rabbits for single-dose (acute toxicity) and repeated-dose (sub-chronic toxicity) showed no abnormalities in the animal’s organs and behaviors and no deaths were reported in tested animals. Two doses of vaccination have been shown to protect NHPs against SARS-CoV-2 infection, as detected by drastic viral reduction from sample swab in nasal, anal, trachea and nasal wash in 7 days after virus challenged, also viral load measurement from lung and BAL tissue showed negative result, which gave better result than negative control and control vaccine group. No evidence of disease enhancement was observed. These results support clinical development of SARS-CoV-2 vaccine, and in 2022 this vaccine has been approved for emergency use in Indonesia.

## Introduction

On March 11^th^, 2020, the World Health Organization (WHO) declared a coronavirus disease 2019 (COVID-19) global pandemic caused by severe acute respiratory syndrome coronavirus 2 (SARS-CoV-2. The disease has impacted global health with substantial cases as well as mortality and severely affected the global economy. The development of a safe and effective vaccine that can be rapidly deployable is an urgent global health prime concern. Numerous vaccine technology platforms and formulations for targeting the SARS-CoV-2 Spike (S) protein, have been pursued and licensed under emergency use authorization (EUA) scheme including nucleic acid vaccines (mRNA and DNA), human and simian replication-defective adenoviral vaccines or viral vector, whole-inactivated SARS-CoV-2, and subunit protein vaccines [1]. Majority of the vaccines have focused on inducing antibody responses against the trimeric SARS-CoV-2 S protein. S protein contains a receptor-binding domain (RBD) that specifically interacts with angiotensin-converting–enzyme 2 (ACE2) receptor and triggers virus–cell-membrane fusion during infection [2, 3]. RBD protein has been an attractive target in COVID-19 vaccine development because of its ability to generate neutralizing antibodies that can inhibit the binding process between SARS-CoV-2 and ACE2 receptor. Using RBD protein as a target antigen may reduce potential unwanted effects such as risks from antibody-dependent enhancement (ADE) since unnecessary part from the S protein is excluded [1, 4].

As of June 10, 2021, 2.2 billion doses of COVID-19 vaccine have been provided, but most of these vaccines have been manufactured and given to people in high-income countries, and the availability of vaccines was delayed for people in low- and middle-income countries (LMICs). The efforts to ensure timely and wider access to vaccines are by facilitating the manufacture of vaccines locally in countries that have public sector vaccine manufacturing [5]. Therefore, the development of a COVID-19 vaccine to address the supply gap in the countries and their regions is imperative.

We developed a COVID-19 vaccine based on yeast-expressed recombinant protein containing SARS-CoV-2 spike RBD protein antigen. This vaccine was developed in collaboration between Bio Farma-Indonesia and the Texas Children’s Hospital Center for Vaccine Development at Baylor College of Medicine (BCM). The antigen was constructed based on sequence of the wild-type SARS-CoV-2 RBD amino acid, representing residues 331-549 of the spike (S) protein (GenBank: QHD43416.1) of the Wuhan-Hu-1 isolate (GenBank: MN908947.3). To enhance immunogenicity of the vaccine, RBD was formulated with Alhdyrogel and CpG1018. Both adjuvants have good safety profiles and have already been used in various licensed vaccines for use in humans such as Alhydrogel is used in Hepatitis B vaccine produced by PT Bio Farma, CpG1018 is used in Heplisav B vaccine produced by Dynavax, and Corbevax produced by Biological E [6, 7]. Nonclinical studies of the COVID-19 vaccines were carried out to evaluate safety in both acute as well as repeated toxicity and the immunogenicity profiles of the COVID-19 vaccine before entering the clinical evaluation stages. The studies include immunological tests that consist of antibody titer and neutralization tests, single-dose for acute toxicity, and repeated-dose for sub-chronic toxicity studies in rodents (Wistar rats) and non-rodents (New Zealand rabbits) animals. After the immunological and safety testing stages have been proven and have good results, this new COVID-19 vaccine candidate was tested for challenge tests on non-human primates (NHP), Macaca fascicularis. The nonclinical studies on receptor binding domain (RBD) COVID-19 vaccine comply with the WHO Guidelines on the nonclinical evaluation of vaccines, WHO Technical Report Series (TRS) 927 Annex 1, and WHO Recommendations to ensure the quality, safety, and efficacy of recombinant Covid-19 vaccines, WHO TRS 978 Annex 4 [8, 9]. Here, we report the preclinical studies in rodents, non-rodents, and non-human primates.

## Materials and methods

### Ethical approval statement

The animal studies, an immunogenicity study in Balb/c mice was conducted by Biofarma. Acute toxicity and repeated dose toxicity study in Wistar rats and New Zealand rabbits were conducted by School of Pharmacy, Bandung Institute of Technology, and the immunogenicity and challenge testing in Non-Human Primates (Macaca fascicularis) were conducted by Professor Nidom Foundation (PNF), Indonesia. The immunogenicity study protocol was approved by the Institutional Animal Care and Use Committee (IACUC) PT Bio Farma (No. 01/IACUC-BF/XI/21). The acute toxicity and repeated dose toxicity study was approved by the Ethics Committee for Animal Research of Bandung Institute of Technology (No. 05/KEPHP-ITB/11-2021; 06/KEPHP-ITB/11-2021; 07/KEPHP-ITB/11-2021; 08/KEPHP-ITB/11-2021). The immunogenicity and challenge testing in Non-Human Primates protocol was approved by the Institutional Animal Care and Use Committee - Professor Nidom Foundation (IACUC-PNF) (No. 2104/IACUC/XII/2021).

### Study design of immunogenicity in BALB/c mice

This study was conducted to determine the immunogenicity profile including antibody titer response neutralization profile of the RBD candidate vaccine with combination of Alhdyrogel and CpG adjuvant in the BALB/c mice. Female BALB/c mice (inbred), aged 6-8 weeks with a body weight of 20 to 25 g (n=20 per group; total 11 groups) imported from Nomura Siam International - Thailand and monitored for 1-5 days in PT. Biofarma’s animal quarantine facilities. They were immunized with a vaccine containing RBD 1.25 μg, 2.5 μg, 5 μg, and 10 μg formulated with Alhydrogel 100 μg alone or a combination of Alhydrogel 100 μg and CpG 20 μg with a total volume of 100 μL by the intramuscular (IM) route on the left and right thighs of mice on day 0 and day 21 [Fig. 1]. The vaccine adjuvant combination Alhydrogel 100 μg + CpG 20 μg and CpG only were used for negative control groups. Control vaccine (inactivated vaccine) was used as a comparison group. Using the submandibular bleeding method on the second day and the heart puncture method on the 35th day, blood samples were obtained to assess the reaction to the first and second vaccinations. The responses observed were specific anti-RBD antibody titer measured by the ELISA method and antibody neutralization assay was done using MNT and the profiles assessed by the surrogate virus neutralization test (sVNT) kit (Genscript).

**Fig 1.**
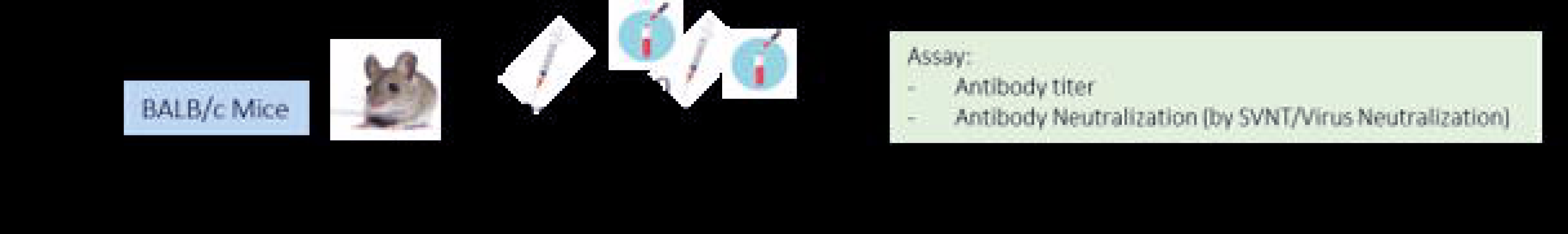
Immunogenicity in BALB/c mice. Time course of immunization and sampling. Mice (n=20) were vaccinated with 1.25μg, 2.5μg, 5μg and 10μgRBD adjuvanted with 100μg Alum only or combination with 20μg CpG 1018, Alhydrogel 100μg and CpG 20μg, RBD 10μg only and control vaccine.

All animal procedures were approved by the Institutional Animal Care and Use Committee (IACUC) PT Bio Farma (No. 01/IACUC-BF/XI/21), and were performed in accordance with all relevant regulatory standards. Mice were housed under 12 h light/dark cycle conditions. Ambient temperature is 22 °C and humidity is 50%. Food and water were provided ad libitum.

### Study design of acute toxicity in Wistar rats and New Zealand rabbits

The rodents used for the toxicity test were male and female Wistar rats aged 6-8 weeks with a body weight of 200-250 grams supplied from Bio Farma Internal Animal Breeding - Cisarua and then monitored and conditioned for 1-5 days in animal quarantine facilities. For non-rodent animals, the toxicity test used male and female New Zealand strain rabbits aged 3-4 months with a body weight of 2.3-2.7 kg supplied from Internal Animal Breeding Bio Farma-Cisarua then monitored and conditioned for 1-5 days in animal quarantine facility.

Single IM dose toxicity of RBD COVID-19 vaccine was evaluated in Wistar rats (20 animals per group: 10 males, 10 females) and New Zealand white rabbits (10 animals per group: 5 males, 5 females). Both were administered with dose levels vaccine of 12.5 and 25 μg/dose with various adjuvant dose combinations. A total of 160 Wistar rats, 80 males and 80 females, with body weights between 200 g and 250 g and 80 New Zealand rabbits, 40 males and 40 females, with body weights between 2.3 kg and 2.7 kg were randomly assigned into 8 dose groups for the study as described in Fig. 2 and Table 1. Up to 0.5 mL of the vaccine was administered intramuscularly to each rat and rabbit in a single dose, and observations were recorded for up to 14 days thereafter. Later, animals were euthanized via cardiac puncture. The behaviour, mortality, clinical symptoms and local reactions, body weights, animal feed consumption, hematology, organ index and tissue histology, urinalysis and biochemistry were observed and recorded for 14 days.

**Fig 2.**
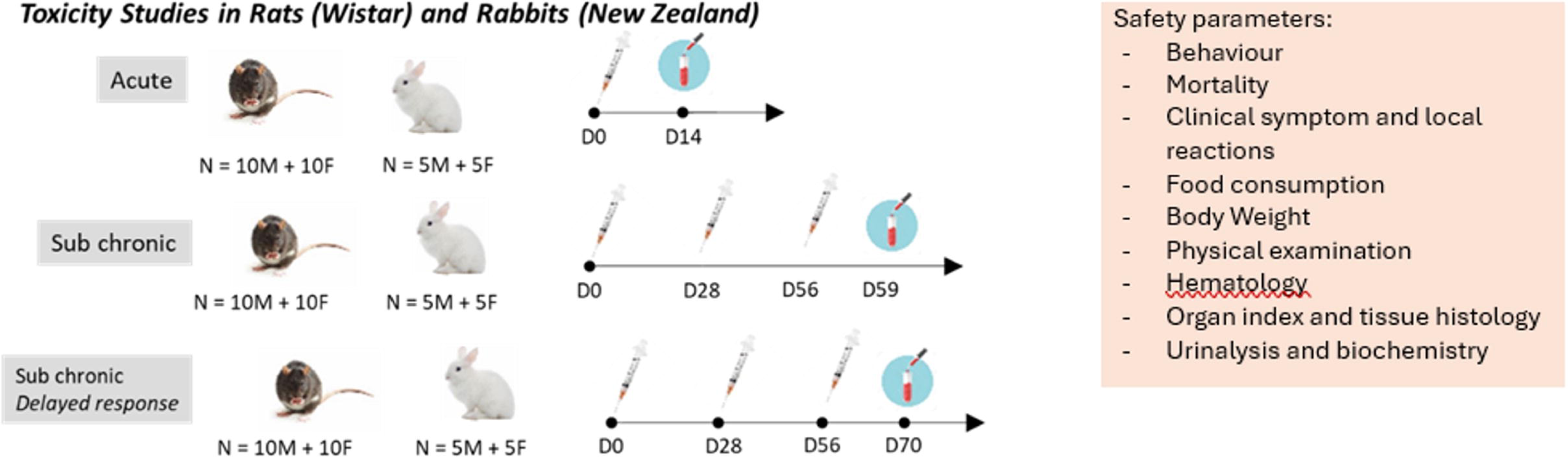
Toxicity studies in Wistar. Time course of immunization and sampling for acute, sub chronic and sub chronic delayed response. Mice (n=20, 10F and 10M) were vaccinated with 12.5μg and 25μg RBD adjuvanted with 750 μg Alum and 750 μg or 2500 μg CpG 1018 μg, Alhydrogel only 750 μg, CpG only 1500μg and combination Alhydrogel and CpG and Saline.

**Table 1.**
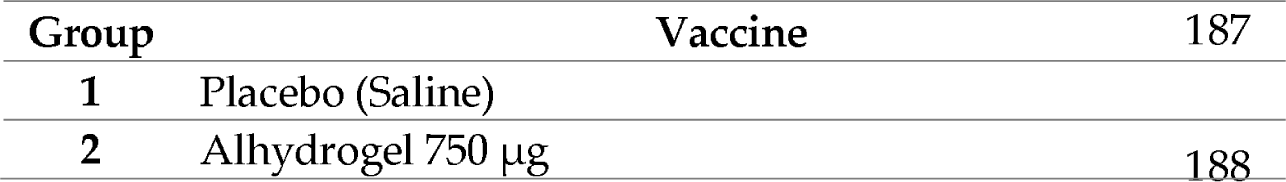
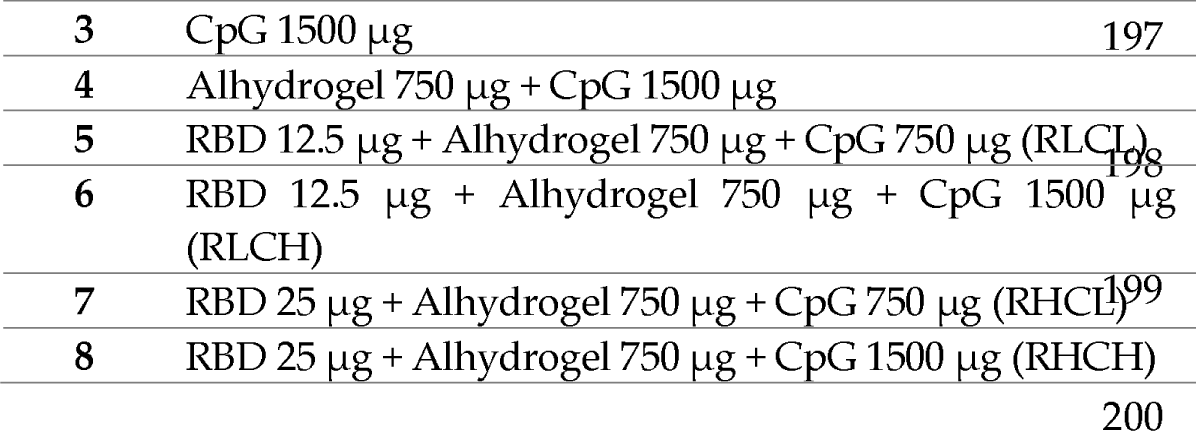
Vaccination dose of toxicity studies in Wistar rat and New Zealand rabbits.

### Study design of repeated dose (sub chronic) toxicity in Wistar rats and New Zealand rabbits

Two repeated-dose toxicity studies were carried out to gain an overview of the cumulative toxicity of a vaccine that has been administered repeatedly by intramuscular route over a predetermined duration of time. Repeated-dose toxicity of RBD Covid-19 vaccine was evaluated in 6–8-week-old Wistar rats (20 animals per group: 10 males, 10 females) and 3-4 months old New Zealand white rabbits (10 animals per group: 5 males, 5 females). Both were administered with dose levels vaccine of 12.5 and 25 μg/dose with various adjuvant dose combinations. To determine the reversibility, persistence, and delayed effect, a satellite group was used wherein the treated animals were further observed for 14 days. A total of 320 Wistar rats with body weights between 200 g and 250 g and 160 New Zealand rabbits with body weights between 2.3 kg and 2.7 kg were randomly assigned into 8 dose groups for the study as described in Fig 2 and Table 1. Up to 0.5 mL of the vaccine was administered intramuscularly to each rat and rabbit on Day 0, Day 28, and Day 56. Testing animals were euthanized on day 59, while satellite animals (sub-chronic delayed response) were euthanized on day 70. The behavior, mortality, clinical symptoms and local reactions, body weights, animal feed consumption, hematology, organ index and tissue histology, urinalysis, and biochemistry were observed and recorded for 70 days.

### Study design of challenge test of Macaca fascicularis

Immunogenicity and challenge tests were conducted on male Macaca fascicularis (3 animals per group), (negative for type D retrovirus, simian immunodeficiency virus, simian lymphocyte tropic virus type 1, and SARS-CoV-2), with an average weight of 3-7 kg or age 2-4 years and supplied from a local Indonesian vendor then monitored and conditioned for 14 days before treatment.

The test group used three Macaca fascicularis in each formulation group (6 formulations, Tabel 2.) with a total of 18 test animals and 2 additional as reserves. The test animals were vaccinated with 0.5 mL intramuscularly on day 0 and day 28. Sera from NHP were taken on days 28, 42, and 56 for the immunogenicity test [Fig. 3]. A challenge study was conducted on 2 variants of the COVID-19 virus. The first batch of challenge tests was infected with <u>hCoV-19/Indonesia/JI-PNF-211352/2021 (SARS-CoV-2 strain Wuhan-Hu-1)</u>. While the second batch of challenge tests was infected with <u>hCoV-19/Indonesia/JI-PNF-213674/2021 (DELTA)</u>. The challenge test was carried out one month after the day 28^th^, which was the day 56^th^ via the intratracheally route and the nostril intranasally route with totally with 2.5 x 10^5^ plaque-forming units (PFU) SARS-CoV-2. The test observations were performed on Macaca fascicularis and recorded until the animals were euthanized are mortality, clinical symptoms, food consumption, body weight, physical examination, hematology and coagulation, blood chemistry, humoral immune and cellular immune response, antibody neutralization, viral loads in swab and tissue, histopathology, immunochemistry and viral component in organ tissue. All procedures were performed under anaesthesia (ketamine, 10 mg/kg body weight, intramuscularly).

**Fig 3.**
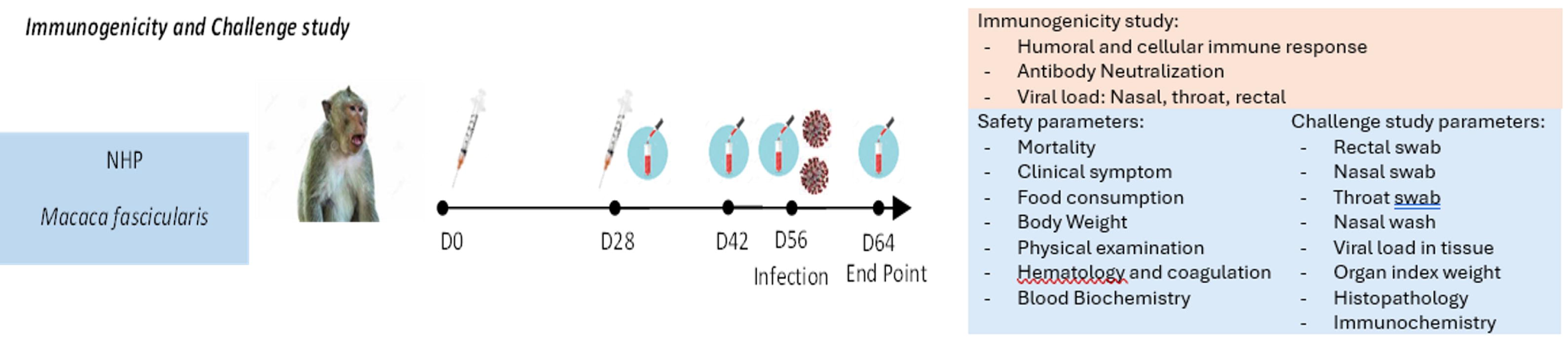
Challenge testing in non-human primates with Wuhan and Delta Coronavirus Indonesian strain. Non-human primates were vaccinated with 12.5μg and 25μg RBD adjuvanted with 750 μg Alum and 750 μg or 2500 μg CpG 1018 μg, Alhydrogel only 750 μg, CpG only 1500 μg and combination Alhydrogel and CpG and Saline and challenged with sars-cov-2 Indonesian strain virus.

**Table 2.** Vaccination dose in Non-Human Primates.

| Group | Vaccine | Number of animals | Route of vaccination |
| --- | --- | --- | --- |
| 1 | RBD 12.5 µg + Alhydrogel 750 µg + CpG 750 µg (RLCL) | 3 | Intramuscular (IM) |
| 2 | RBD 12.5 µg + Alhydrogel 750 µg + CpG 1500 µg (RLCH) | 3 |  |
| 3 | RBD 25 µg + Alhydrogel 750 µg + CpG 750 µg (RHCL) | 3 |  |
| 4 | RBD 25 µg + Alhydrogel 750 µg + CpG 1500 µg (RHCH) | 3 |  |
| 5 | Vaccine control | 3 |  |
| 6 | Placebo (Saline) | 3 |  |

### Humoral immune response (ELISA)

To assess the immune response, serum samples were collected from immunized animals after the first vaccination (day 20), and after the second vaccination (day 35). ELISA plates (Maxisorp, Nunc-Immuno plate) were coated overnight at room temperature with 5 μg/mL of RBD protein in phosphate buffer saline and blocked with 10 g/mL BSA in PBS. Diluted sera were applied to each well for serum samples overnight at room temperature. The next day plates were incubated with goat anti-mouse IgG-HRP antibody and subsequently developed with 3,3’,5,5’-tetramethylbenzidine (TMB) substrate. Reactions were stopped with 2M sulfuric acid, and the absorbance was measured at 450 nm using a microplate reader. For macaques, serum samples were collected on days 28, 42, 56, and 63 days. Before analysis, serum samples were inactivated at 56^0^C for one hour. ELISA plates were coated overnight at 4^0^C with 2 μg/ml of RBD protein and blocked in 2% skim milk in PBS. Serum samples were serially diluted and added to each well. Plates were incubated with goat anti-monkey IgG-HRP antibody and developed with 3,3’,5,5’-tetramethylbenzidine (TMB) substrate. Reactions were stopped with 2 M sulfuric acid, and the absorbance was measured at 450 nm using a microplate reader [10]. The antibody titers were determined by endpoint dilution and calculated using GraphPad Prism version 8.4.3 for Windows (GraphPad Software). For mice, cut off (baseline) value was defined from negative control sera of alhydrogel-CPG combination (without RBD) and RBD only without adjuvant [11]. Cutt-off/baseline values from antibody titer macaques were defined experimentally and depended on assay background and noise. When the OD450 value of the serum tested ≤ cut off value, the endpoint titer was zero.

### Neutralizing assay

There are several serological assays to detect neutralizing antibodies, Micro Neutralization Test (MNT) and Plaque Reduction Neutralization Test (PRNT) require handling live viruses in biosafety level 3 (BSL 3) containment facility. MNT assay is similar to a PRNT, but it can be performed in a 96-well cell culture plate and allows for a higher throughput compared to a standard PRNT [12]. Surrogate virus neutralization test (sVNT) is a method to detect neutralizing antibodies without the need for any live virus or cells, and can be completed in 1-2 hours in BSA2 laboratory [13]. In this study, we performed sVNT and MNT to detect neutralizing antibodies in mice sera, and PRNT for NHP sera. MNT assay was provided by National Institute of Health Research and Development Republic of Indonesia (NIHRD). The viruses used for MNT and PRNT are SARS-CoV-2 Wuhan strain and Delta strain. The cell line used was Vero cells. Naive sera were used as negative control, while neutralizing mAb or antiserum was used as positive control. Both MNT and PRNT methods can refer to the Bewley method [12]. Surrogate VNT reagents and assay method were prepared according to the Tan C. W. et al method [14].

### Cellular immune response

To assess the cellular immune response, ELISPOT assay was performed using ELISpot Plus: Monkey IFN-γ, IL-4, IL-5, IL-6, and tumor necrosis factor-alpha (TNF-α) Kits (MabTech). Pre-coated plates were conditioned with medium containing 10% of the same serum as used for the cell suspension. Peripheral Blood Mononuclear Cells (PBMC) were incubated with stimuli, peptide pool SARS-CoV-2 200µl/well for 12-24 h, 37^0^C with 5% CO_2_. Plates were incubated with detection antibodies for Interferon-gamma (IFN-γ), IL-4 and IL-5, IL-6 and tumor necrosis factor-alpha (TNF-α). Incubation plate continued with streptavidine-HRP and developed with TMB substrate solution. When the colored spots were intense enough to be observed, the development was stopped by thoroughly rinsing samples with deionized water. The numbers of the spots were determined using an automatic ELISPOT reader and image analysis software.

### Viral loads in swab and tissue

Sample tissue and swabs were soaked and grinded in saline. Virus genomic RNA was isolated from supernatant of grinded tissue and soaked swabs using Simply P Virus DNA/RNA Extraction Kit Cat No. BSC67M1 (Bioflux). Quantitative reverse transcription-PCR (qRT-PCR) assays were tested with SuperScript^TM^ III Platinum^TM^ One-Step qRT-PCR Kit (Invitrogen) on Quant Studio 5, AB according to manufacturer’s protocol. Primers and probes from 2019-nCoV RUO Kit, IDT, were used to detect Nq and N2 regions of viral genome.

### Organ index and histopathology analysis

For organ index examination, isolated animal organs were washed in saline water, drained, and weighed. The organs are brain, liver, lungs, spleen, kidney, heart, ovary, and uterus (female rat), and testis (male rat). For histopathology analysis, animal organs were fixed in 10% neutral buffer formalin for 24h, soaked in ethanol and xylol, embedded in liquid paraffin, and then sectioned. Tissue section, 5 µm, was deparaffinized in xylene and stained with hematoxylin and eosin (HE). Pathological examination were carried out using Nikon Eclipse Ci, NIS-Element D4.5.0064-bit programme with 400x magnification. Evaluation of histopathology was scored based on the Klopflrish method [15].

### Immunochemistry in organ tissue

The paraffin tissue sections were deparaffinized with xylene, rehydrated through successive bathes of water, and incubated in 3% H_2_O_2_ at room temperature. Subsequently, the sections were blocked with sniper, washed with PBS, incubated with primary antibody against S1 gene SARS-CoV-2 (Gene Tex), and then incubated with Tekavidin-HRP label. The section was counterstained with hematoxylin, rehydrated, and cleaned using xylene.

### Hematology analysis

Blood from sacrificed rats were collected by heart puncture method. Hematology analysis was performed using hematology analyzer.

### Blood biochemical observation

To assess the blood biochemical observation, serum samples were collected from immunized animals. Serum was tested with chemical reagents to measure urea, creatinine, albumin, alkaline phosphatase (ALP), calcium, glucose, and serum glutamate-pyruvate transaminase (SGPT).

### Urine profile

Urin profile was observed for the pH value and specific gravity.

### Observation of animal body weight and feed consumption

For animal body weight, every animal was weighed every morning based on schedule.

## Results

### Immunogenicity evaluation in BALB/c mice

Antibody response in mice that received RBD adjuvanted with Alhydrogel with RBD doses of 1.25µg, 2.5µg, 5µg, or 10µg, after one administration increased slightly compared to negative control (adjuvant - only). After two administrations, antibody titers were increased with the two highest titers (±SD) were 3.40 (±0.97) and 3.14 (±0.86) for the group of mice receiving RBD doses of 2.5µg and 5µg respectively. When mice were vaccinated with the RBD adjuvanted with a combination of Alhydrogel and CpG, the titer was significantly higher after receiving one and two administrations with the two highest titers (±SD) were 4.92 (±0.34) and 4.84 (±0.4) post first administration and 5.83 (±0.37) and 5.91 (±0.31) after 2^nd^ administration for RBD doses of 2.5µg and 5µg respectively. The mice group injected with vaccine control (inactive vaccine) had antibody titer 3.88 (±0.22) after the first injection and 5.07 (± 0.26) after the second injection. If we compare to the adjuvant groups, the addition of Alhydrogel adjuvant increased the titer by 1.57 – 1.85-fold compared to the baseline titer of adjuvant negative control without RBD, while adjuvant combination of Alhydrogel and CpG increased the average titer by 3.1 – 3.21-fold compared to the group adjuvant without RBD. On the other side, vaccine control has a 2.75-fold antibody titer compared to adjuvant control. The details of the titer antibody data are provided in Table 3.

**Table 3.** Antibody titer after one and two vaccine administration.

| No. | Group | Antibody titer after one<br>vaccine administration<br>(mean ± SD) | Antibody titer after two<br>vaccine administration<br>(mean ± SD) |
| --- | --- | --- | --- |
| 1 | Alhydrogel+CpG | 1.65 ± 0.15 | 1.84 ± 0.27 |
| 2 | RBD 10 μg | 1.70 ± 0.31 | 2.26 ± 0.44 |
| 3 | RBD 1.25 μg + Alhydrogel | 2.00 ± 0.12 | 2.95 ± 0.84 |
| 4 | RBD 2.5 μg + Alhydrogel | 1.99 ± 0.24 | 3.40 ± 0.97 |
| 5 | RBD 5 μg + Alhydrogel | 1.98 ± 0.10 | 3.14 ± 0.86 |
| 6 | RBD 10 μg + Alhydrogel | 2.08 ± 0.20 | 2.90 ± 0.89 |
| 7 | RBD 1.25 μg + Alhydrogel +<br>CpG | 4.41 ± 0.66 | 5.73 ± 0.33 |
| 8 | RBD 2.5 μg + Alhydrogel +<br>CpG | 4.92 ± 0.34 | 5.83 ± 0.37 |
| 9 | RBD 5 μg + Alhydrogel + CpG | 4.84 ± 0.40 | 5.91 ± 0.31 |
| 10 | RBD 10 μg + Alhydrogel + CpG | 4.62 ± 0.83 | 5.76 ± 0.39 |
| 11 | Control vaccine (1/5 shd) | 3.88 ± 0.22 | 5.07 ± 0.26 |

When comparing the responses in group vaccine control and RBD adjuvanted with Alhydrogel-CpG after two administrations, it was found that antibody response from the RBD vaccine was significantly higher compared to the control vaccine [Fig. 4a]. On the other hand, there was no noticeable difference between the various RBD doses tested in this study. The antibody neutralization profile was determined from the second post-administration, sera obtained from each sample of mice and determined with the sVNT and MNT methods. sVNT method using kit for Wuhan strain, and neutralization antibody profiles for animals that received the RBD formula with the adjuvant combination Alhydrogel and CpG were confirmed with microneutralization (MNT) method using Wuhan and Delta Strain. In the neutralization profile of the negative control group, sera had a low neutralization ability, while the antibodies from the group of mice vaccinated with the RBD-Alhydrogel adjuvanted vaccine showed a slightly higher neutralization response and for neutralization titer produced by the RBD vaccine with the adjuvant combination of Alhydrogel and CpG was significantly higher than negative control and RBD-Alhydrogel group, and it is equivalent to control vaccine [Fig. 4b]. For the MNT neutralization test variant against the Wuhan and Delta variants [Fig. 4c], the trend of neutralization titer was decreased between the effects of neutralization of the Wuhan compared to Delta variant. The antibody titer against Wuhan compared to Delta varian was decreased by 0.52; 0.17; 0.1; 0.35 and 0.88 fold for mice sera were vaccinated with RBD 1.25µg, 2.5µg, 5µg, and 10µg with combination of Alhydrogel and CpG adjuvant. The neutralization titer of RBD with Alhydrogel and CpG adjuvant was higher than the control vaccine [Fig. 4c]. The 5 µg and 10 µg RBD dosages and 2,5 µg and 5 µg RBD dosages were shown to have the highest response respectively against Wuhan and Delta when compared to other doses.

**Fig 4.**
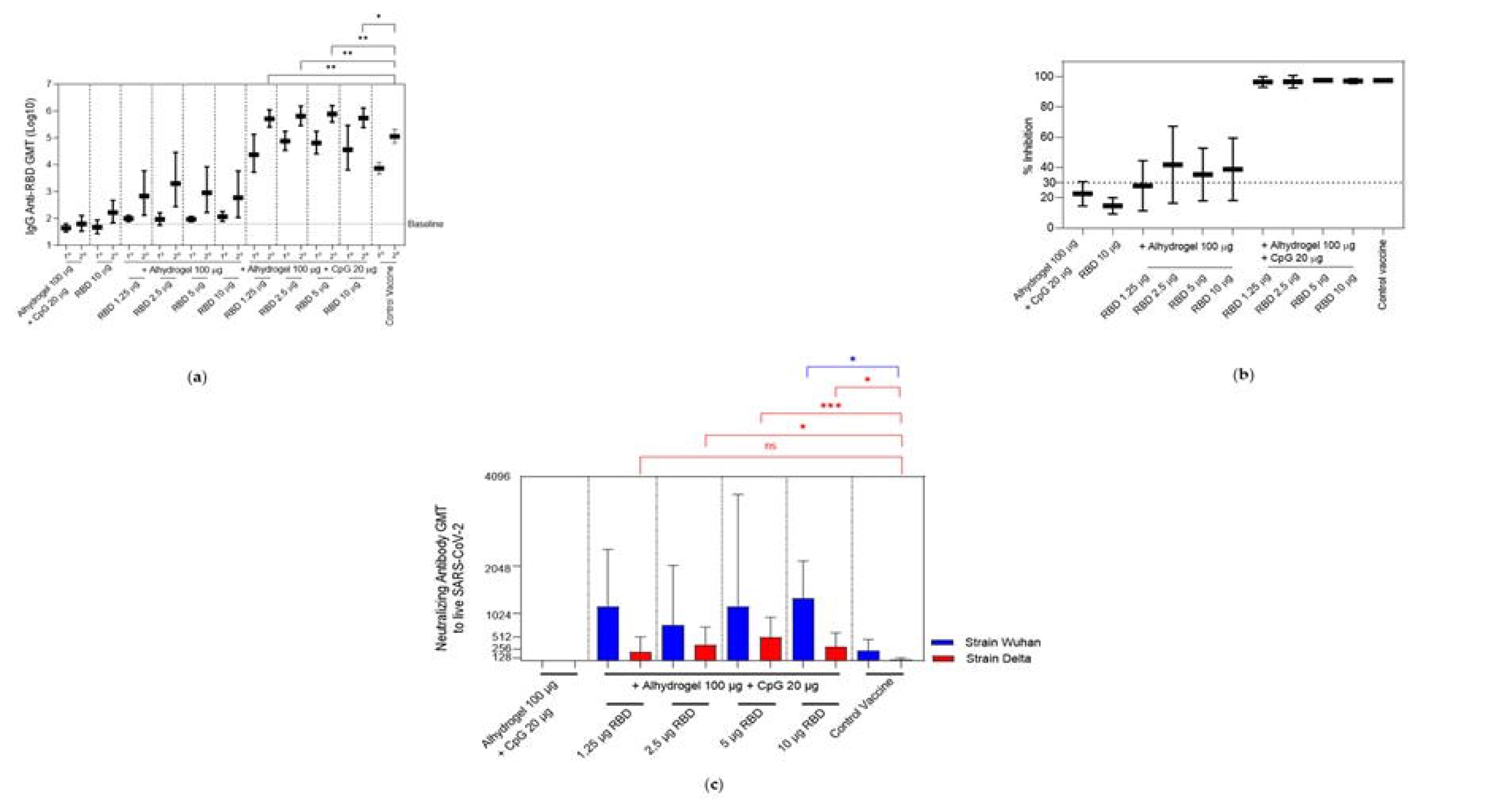
Immunogenicity in BALB/c mice. **(a)** Antibody titer after first (1°) and second (2°) vaccine administrations. The negative control group or baseline was from the group vaccinated with a combination Alhydrogel+CpG and RBD only. The group that was vaccinated with control vaccine was used as a comparison group. Sera from mice were taken on the 20^th^ and the 35^th^ days. Antibody titer data were analyzed with an ordinary one-way ANOVA followed by Turkey’s multiple comparisons test (**p<0.01, *p<0.05) that showed significant differences. (**b**) sVNT result for SARS-CoV-2 (Wuhan strain) neutralization assay. The data showed % neutralization result between negative control, RBD adjuvanted Alhydrogel, RBD adjuvanted Alhydrogel+CpG, and control vaccine group. (**c**) MNT result for antibody neutralization profile to Delta and Wuhan strain. Data values were analyzed with an ordinary one-way ANOVA followed by Tukey’s multiple comparisons test (ns>0.05, *p<0.05, ***p<0.001) that showed significant differences. The mean values presented with standard deviation (SD).

### Acute toxicity test in Wistar and New Zealand rabbits

#### Observation of clinical symptoms and local reactions

No significant difference in clinical symptoms and local reactions at the injection site in all test groups was observed until the 14^th^ day after administration of the test preparation compared to the control group. None of the animals died during the observation period.

#### Observation of animal body weight and feed consumption

The vaccinated rats (both male and female) from all groups had an increase in body weight during the observation period when compared to the control group, the animals in the test group had a higher trend of increase in weight gain. Female rabbits showed an increase in body weight during the observation period following vaccination, which was similar between the control and other groups. The increase in body weight was within the normal range. In male rabbits, the profile of changes in body weight in each group varied. Male rabbits in the normal control group and Alhydrogel (Alum) 750 µg + CpG 1500 µg showed a decrease in body weight while the other six groups showed an increase in body weight. Neither Wistar rats nor New Zealand rabbits showed no significant difference in the amount of feed consumed by test animals until the 14^th^ day of the observation period.

#### Hematological analysis

Hematological parameters tested during the study were white blood cell (WBC), hemoglobin (HGB), mean corpuscular hemoglobin (MCH), mean corpuscular hemoglobin concentration (MCHC), red blood cell (RBC), mean corpuscular volume (MCV), hematocrit (HCT), and platelets (PLT). No significant difference in WBC, HGB, MCH, MCHC, RBC, MCV and HCT result from both rats and rabbits from all groups. In rats, results showed that the PLT count was significantly lower (P<0.05) in the CpG 1500 µg and Alum 750 µg + CpG 1500 µg groups, as well as inconsistent changes in the RBD 12.5 µg and 25 µg groups as compared to the control group. However, most of these changes in values remained within the range of platelet values in the control group, which were 150-309 x10^3^/mL for male rats and 120-247 x10^3^/mL for female rats. In female rabbits, PLT levels were significantly lower (P<0.05) on Day 14 in the normal control group and RLCL groups compared to Day 0 (before the administration of the test preparation. In male rabbits, PLT levels were lower in most of the test groups on day 14 but significantly lower (P<0.05) in the RLCL groups compared to Day 0 (before the administration of the test preparation).

#### Observation of organ index and tissue histology

In organ index examination, the results showed that the liver index of male and female rats in the RLCL group was significantly lower (P<0.05) than in the control group. However, in the other groups, RHCH, RHCL, RLCH showed significant differences compared to the control group. In female rabbit organs showed that the liver index was significantly lower (P<0.05) in the CpG 1500 µg group, Alhydrogel 750 µg + CpG 1500 µg, RHCL, RHCH, and RLCH compared to the normal control group. The liver index of male rabbits in the Alhydrogel 750 µg and RLCL groups was significantly lower (P<0.05) than the normal control group.

Liver safety can be further confirmed by histopathological observations of the liver. In the histological observation of liver tissue, the shape of the vascular cavity and the distribution of Kupffer cells were similar in the normal control group (saline) and the test groups. Histological observations of other organs (brain, lungs, heart, kidney, thymus, spleen, and testes) were similar between the normal control group and the test groups.

#### Urine profile

In urinalysis, no significant changes were observed in the pH value and specific gravity of the urine in all the test groups. The pH values and urine-specific gravity of all test groups were within the normal range.

#### Blood biochemical observation

Blood biochemistry tested during the study were urea, creatinine, albumin, alkaline phosphatase (ALP), calcium, glucose, and serum glutamate-pyruvate transaminase (SGPT). Results showed that ALP levels were significantly higher (P<0.05) in male and female rats in the following groups: RHCH, RLCL, and RLCH. Significantly higher ALP levels in Alhydrogel 750 µg + CpG 1500 µg and RHCL groups were only found in male rats. In female rabbits, all test groups showed significantly higher (P<0.05) glucose levels than the normal control group. According to the literature, normal glucose levels in female rabbits are 89.0-149.9 mg/dL [16]. In male rabbits, there was no difference in the glucose levels of the test animals. In male rabbits, ALP levels were significantly lower (P<0.05) in the Alhydrogel 750 µg + CpG 1500 µg and RLCL groups as compared to the normal control group. However, ALP levels were still in the normal range of 100 - 400 U/L [12]. In male rabbits, the RLCH group showed significantly higher (P<0.05) levels of urea than the normal control group. According to the literature, normal urea levels in male rabbits are 44.1-105.9 mg/dL [16].

### Repeated dose (sub chronic) toxicity in Wistar rats and New Zealand rabbits

#### Observation of clinical symptoms and local reactions

No significant difference in clinical symptoms and local reactions at the injection site in all test groups was observed until the end of the observation period. None of the animals died during the observation period.

#### Observation of animal body weight and feed consumption

The vaccinated animals (both male and female rats and rabbits) from all groups had an increase in body weight after vaccination during the observation period until the last day of necropsy.

#### Hematological analysis

Hematological parameters tested during the study were white blood cell (WBC), hemoglobin (HGB), mean corpuscular hemoglobin (MCH), mean corpuscular hemoglobin concentration (MCHC), red blood cell (RBC), mean corpuscular volume (MCV), hematocrit (HCT), and platelets (PLT). No significant difference in WBC, HGB, MCH, MCHC, RBC, MCV, and HCT results from both rats and rabbits from all groups. In rats, there was a significant difference (P<0.05) observed in the PLT values which were inconsistently observed in the test and satellite groups. In female rabbits, PLT levels in the RHCH groups were significantly lower (P<0.05) than in the normal control group; however, the PLT levels were within the normal range. In the female rabbit satellite group, PLT values were significantly decreased (P<0.05) in the RLCH and RLCH groups than in the normal control group. In male rabbits, PLT levels in all the groups except RLCH group were significantly lower (P<0.05) than in the normal control group; however, the PLT levels were within the normal range. In the male rabbit satellite group, there were no significant differences in the PLT values among the group except for the Alum + CpG group which had a significantly lower (P<0.05) PLT value than the normal control group.

#### Observation of organ index and tissue histology

The liver index of male rats in the Alum 750 µg group was significantly lower (P<0.05) than in the control group. The liver index of male rats in the CpG 1500 µg, Alhydrogel 750 µg + CpG 1500 µg, RLCL, RLCH, and RHCH groups were significantly higher (P<0.05) as compared to the control group. Similar trends were observed for female rats in the main group. In general, there were no significant differences in most organs in all test and satellite groups, including the adjuvant group. In female rabbits, the liver index value in the CpG, Alum + CpG, and RHCH groups showed significantly lower (P<0.05) than the normal control group, while the RHCL group showed significantly higher (P<0.05) liver index value than the normal control group. In male rabbits, the liver index value in the RHCL group showed a significantly higher (P<0.05) value than the normal control group. In the female rabbit satellite, the liver index in the CpG, RLCL, and RHCH groups were significantly lower (P<0.05) than in the normal control group. In the RLCH satellite group, the liver index value was higher than the normal control satellite but not significant (p>0.05). In the male rabbit satellite, the liver index value in the CpG and Alum + CpG groups showed significantly lower (P<0.05) than the normal control group.

#### Blood chemistry analysis

Biochemistry parameters tested during the study were urea, creatinine, albumin, ALP, calcium, glucose, and SGPT. In rats, the analysis showed that there was a significant difference (P<0.05) in the glucose levels of the RBD 25 and 12.5 µg groups which were observed inconsistently in the test group. In the satellite group, nearly all groups had blood glucose levels within the normal range, except for the saline group and RLCH group. Similarly, significant changes were seen in the ALP levels from many satellite groups when compared to the saline group, however, these differences were not seen in the 59-day test group. In male and female rabbits, the results of all biochemical parameters (except glucose and ALP) were comparable to the normal control group. In female rabbits, the glucose levels were significantly lower (P<0.05) in Alum and RHCH groups while it is significantly higher (P<0.05) in the CpG group as compared to the normal control group. In the female rabbits satellite, the RLCH and the RHCL satellite groups showed significantly higher (p<0.05) ALP values than the normal control satellite. The glucose levels in the CpG group were significantly higher (p<0.05), and the glucose levels in the RLCH satellite group were significantly lower (p<0.05) than in normal control satellites. In the male rabbit satellite, the glucose levels in the Alum group, Alum + CpG, RLCL, RHCL, and RHCH groups were significantly lower (P<0.05) than in the normal control group. This indicates that the low glucose level is not caused by the administration of the test vaccine and does not depend on the dose of the test vaccine administered.

#### Urine profile

No significant changes were observed in the pH value, urine volume, and specific gravity of the urine in all the test groups. The pH values and urine-specific gravity of all test groups were within the normal range.

#### Immunogenicity and challenge testing in non-human primates with Wuhan and Delta Coronavirus Indonesia strain

The challenge test studies were conducted according to the World Health Organization (WHO) Guidelines on the nonclinical evaluation of vaccine adjuvants and adjuvanted vaccines [17] as well as available literature on the challenge test on NHP [18]. The viruses used in the challenge test are <u>hCoV-19/Indonesia/JI-PNF-211352/2021</u> <u>(SARS-CoV-2 strain Wuhan-Hu-1) and hCoV-19/Indonesia/JI-PNF-213674/2021 (DELTA)</u>. No mortality was observed in experimental animals, Macaca fascicularis after receiving the first and second vaccinations and being infected with any of the SARS CoV-2 variants (Wuhan/Delta virus).

#### Observation of animal body weight, feed consumption and clinical symptoms

The test animals challenged with any of the SARS CoV-2 variants (Wuhan/Delta virus) strain did not experience significant differences between treatment groups in clinical symptoms, local reactions at the injection site, changes in body weight, feed consumption, physical examinations, mucosal conditions, reflex responses, and fecal conditions.

#### Hematology analysis and blood biochemical observation

For hematological, coagulation, and biochemistry parameters in the test animals challenged with any of the SARS CoV-2 virus variants (Wuhan/Delta virus) strain, there was no difference between treatments in all formula groups; the hematology parameters were within the normal range.

#### Humoral immune response (ELISA)

The immunogenicity test showed four groups of animals with vaccine formulas (RLCL, RLCH, RHCL, and RHCH). After two times vaccinations, RBD vaccine with the adjuvant combination of Alhydrogel and CpG induced NHP to produce antigen-specific IgG-binding antibodies. The titer significantly increased (p<0.01, p<0.001, p<0.0001) from the first vaccine administration until day 56 (Fig. 5A.) even though there was a decrease in IgG titers on day 42 and significantly increased again for RLCL, RHCH and control vaccine on day 56. On day 56, there is a significant difference (p<0.001) in the antibody titer between the control vaccine and RLCL and RLCH, although the antibody titers for RLCH and RHCL were comparable to those of the control vaccination.

**Fig 5.**
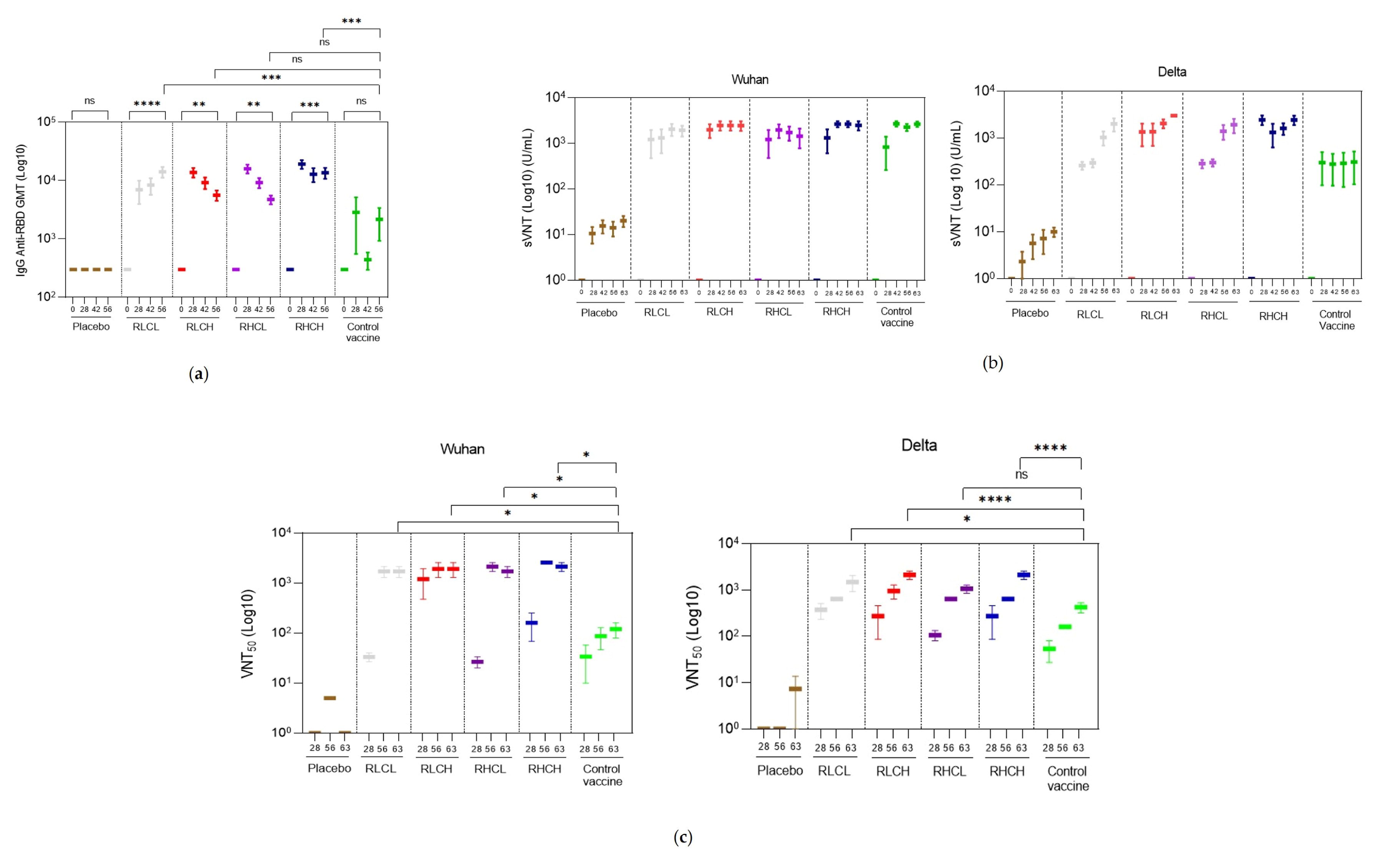
Immunogenicity testing results in non-human primates (Macaca fascicularis). **(a)** Antibody titer after first and second vaccine administrations, and before challenge. The group that was vaccinated with the control vaccine was used as a comparison group. **(b)** Result of Surrogate Virus Neutralization Test (sVNT) to Wuhan-like and Delta strain. The data showed neutralization results after first and second vaccination, and after challenge. **(c)** Result of Plaque Reduction Neutralization Test (PRNT) to Wuhan-like and Delta strain. The data showed neutralization after the first and second vaccination, and after the challenge. The significant differences were determined using two-way ANOVA followed by Turkey multiple comparison test (ns>0.05, *p<0.05, ***p<0.001) that showed significant differences. The mean values are presented with standard errors (SEM).

#### Neutralizing assay

In sVNT result, there was a significant increase in antibody neutralization value after vaccination until day 56 (Fig. 5B). Before day 56 and after infection (day 63), the antibody neutralization value did not show a significant difference. The sVNT result was aligned with PRNT, the vaccine formula (RLCL, RLCH, RHCL, and RHCH) generally showed a significant (p<0.05; ****p<0.0001) high titer of neutralizing antibody in PRNT starting from day 56 (Fig.5C) compared with the control vaccine group which showed a lower degree of neutralization PRNT titer.

#### Cellular immune response

In Wuhan-like observation groups, interferon gamma, showing differences between time and between groups. Groups RHCH, vaccine control, and placebo showed the the highest levels on day 56 and day 63. InterLeukin-4 (IL-4) in this challenge test showed a noticeable difference. IL-4 plays a role in inhibiting pro-inflammatory cytokines and works antagonistically with pro-inflammatory cytokines such as IFN-γ and TNF-α. In this test, IL-4 levels were inversely proportional to IFN-γ, that group RLCL, RLCH, RHCL had higher levels than groups RHCH, control vaccine, and placebo, although the difference was not significant (fig 6).

**Fig 6.**
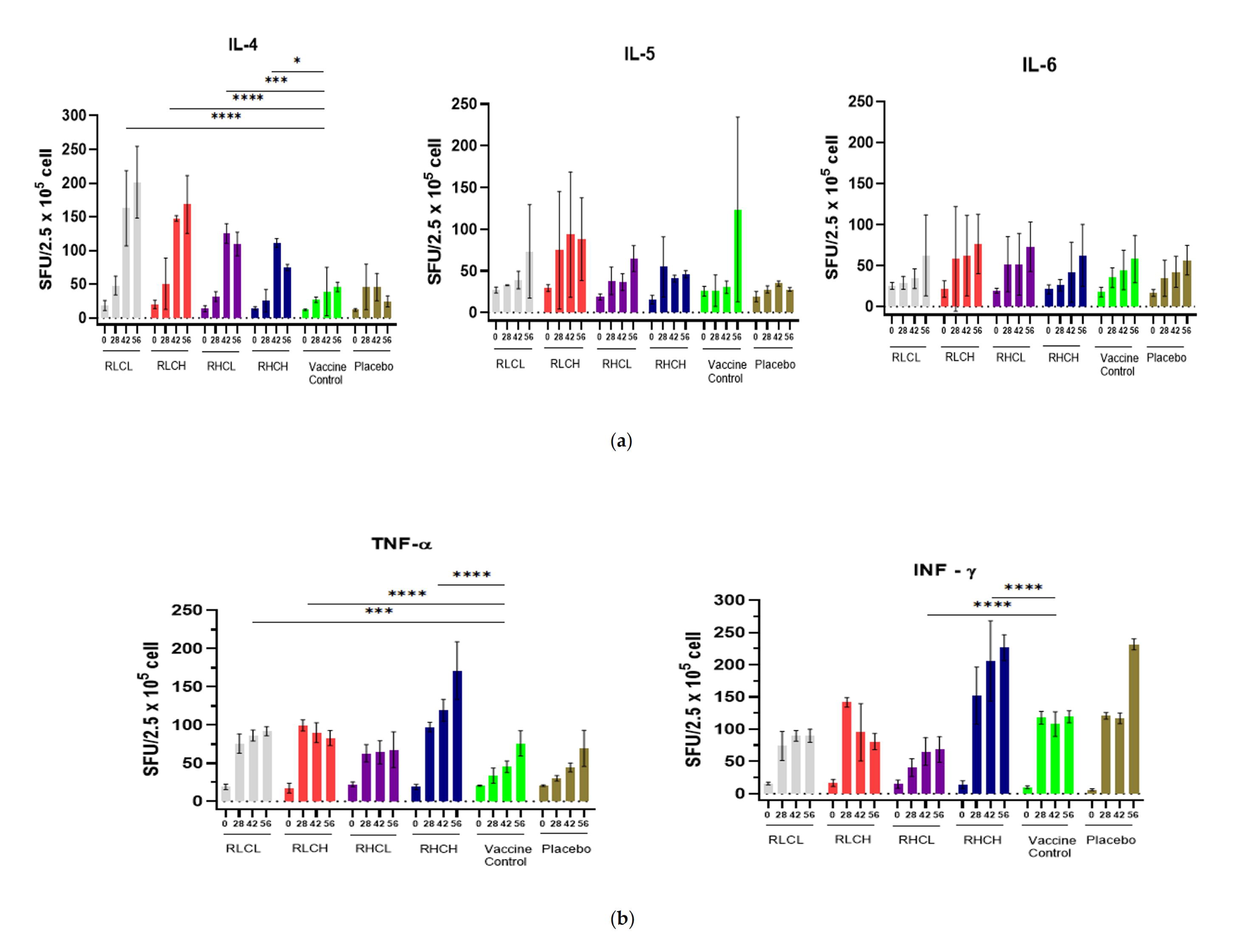
Cellular immune response in non-human primates (NHP) after vaccination and challenge with Wuhan-like strain. Macaques were immunized two times at day 0 and 28 through intramuscular route with RLCL, RLCH, RHCL, RHCH, Placebo, and Control Vaccine. NHP was infected with <u>hCoV-19/Indonesia/JI-PNF-211352/2021</u> <u>(SARS-CoV-2 strain Wuhan-Hu-1)</u> at day 56. (a) Result of IL-4, IL-5 and IL-6; (b) Result of IFN-γ, and TNF-α. The significant differences were determined using two-way ANOVA followed by Dunnett multiple comparison test (*p<0.05; **p<0.01; ***p<0.001; ****p<0.0001).

InterLeukin-5, is a primary cytokine involved in the formation and maturation of eosinophils in the bone marrow and for the passage of eosinophils to the site of infection. In post-vaccination COVID-19 conditions, the animals observed had a lower number of IL-5 than the infected group. In the Challenge test, IL-5 levels in groups RLCL and RLCH were lower than control vaccine and placebo groups. InterLeukin-6 (IL-6) is a pro-inflammatory cytokine that is responsible for host defense and tissue injury. IL-6 will be expressed in a longer period than other pro-inflammatory cytokines. IL-6 can exhibit pathogenic effects, so caution is required when analyzing IL-6 in animal models. In this test, the levels of group vaccine candidates with the CpG formula have a lower infection rate than the comparison control vaccine and placebo. Tumor Necrosis Factor (TNF-α) in this test has a significant effect. TNF-α is a pro-inflammatory cytokine that is a marker of disease severity. In the vaccination group, TNF-α is a precursor to an efficient vaccine response and can provide a predictive marker for a humoral response to vaccination. In general, TNF-α levels will be lower than the infection group. The results of measurements of TNF – α levels showed higher levels in formulas RLCL and RHCH. However, when observing the body temperature of the test animals, no noticeable difference was found, or the body temperature was above normal/fever. In Delta observation groups, interferon gamma in all groups peaked on day 42, and then decreased except for vaccine A which increased again on day 63. For IL-4, IL-5 and IL-6 all groups did not differ significantly at day 63. TNF alpha values on days 28, 42, 56 and 63 did not differ significantly between group sample vaccines but groups control vaccine and placebo had low TNF alpha. In CpG adjuvant vaccines, target cells are dendritic cells which will recognize CpG through TLR 9, and then dendritic cells will secrete IL-6 which will encourage B cell proliferation in the humoral immune system and TNF alpha which will ultimately increase interferon gamma, which in turn leads to a cellular immune response. While the IL-4 and IL-5 examination is to see an allergic response (fig 7).

**Fig 7.**
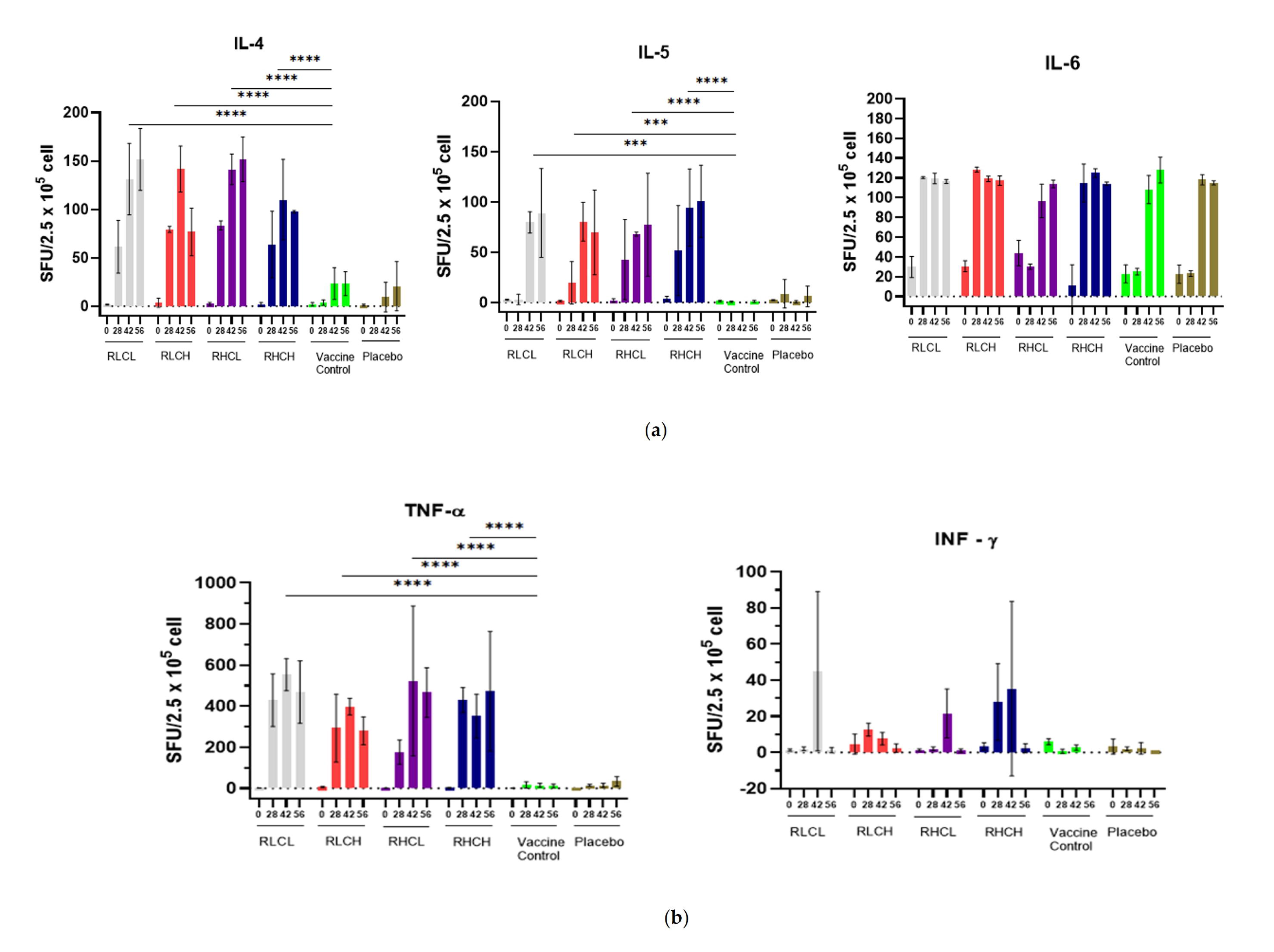
Cellular immune response in non-human primates (NHP). Macaques were immunized two times at day 0 and 28 through an intramuscular route with RLCL (A), RLCH (B), RHCL (C), RHCH (D), Placebo (E), and Control Vaccine (F). NHP was infected with <u>hCoV-19/Indonesia/JI-PNF-213674/2021 (DELTA)</u> at day 56. (**a**) Result of IL-4, IL-5 and IL-6; (**b**) Result of IFN-γ, and TNF-α. The significant differences were determined using two-way ANOVA followed by Dunnett multiple comparison test (*p<0.05; **p<0.01; ***p<0.001; ****p<0.0001).

#### Viral loads in swab and tissue

The results of the viral load measurement from lung and BAL tissue showed negative results for the four test vaccines, which gave better results than placebo and control vaccine (Fig. 8A). Furthermore, the viral load measurements (Fig. 8B) from the sample swab in nasal, anal, trachea and nasal wash showed drastic reduction for the four test vaccines (RLCL, RLCH, RHCL, and RHCH) in 7 days after challenged with Wuhan and Delta virus.

**Fig 8.**
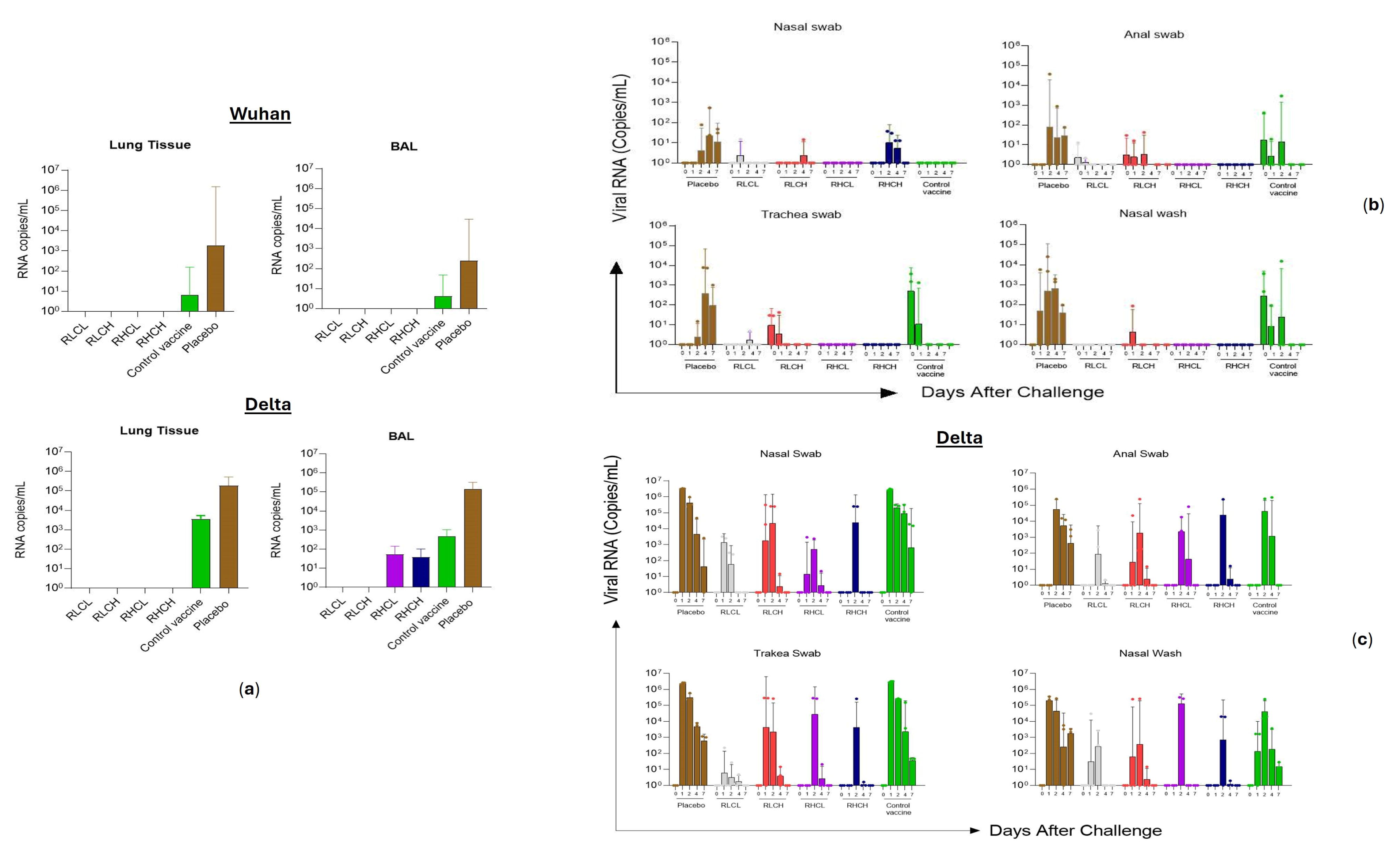
Viral loads in non-human primates (NHP) challenge testing. **(a)** Viral loads in lung tissue and BAL (bronchoalveolar lavage); (**b**) Viral loads Swabs. Samples were taken 7 days after challenge using Wuhan-like virus Indonesia strain; (**c**) Viral loads Swabs. Samples were taken 7 days after challenge using Delta virus Indonesian strain.

#### Immunochemistry in organ tissue

Immunohistochemistry was performed to detect the viral antigens 7 days after challenge in lung tissue, the result showed no viral antigen detected in RBD-containing vaccines when challenged with Wuhan and Delta strain, and the viral antigen was shown in placebo and control vaccine (Fig. 9).

**Fig 9.**
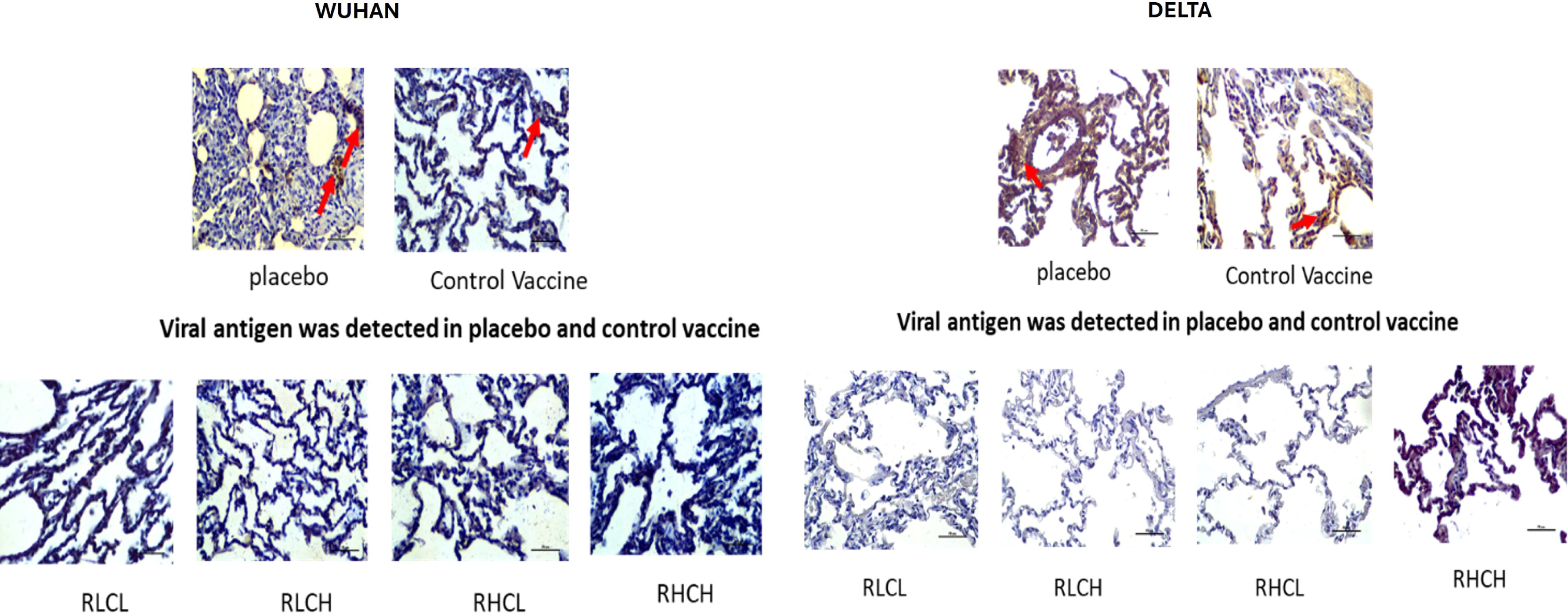
Immunochemistry in organ tissue. Viral component detection test in lung tissue at day 7 post challenged.

#### Histopathology and viral component in organ tissue

Histopathology observations showed that no abnormalities were detected in lung tissue, while infiltration and hemorrhage were detected in placebo and control vaccine (Fig. 10). Likewise in immunochemistry test showed that viral antigen was not detected for the four test vaccines. Whereas some viral antigen was detected in placebo and control vaccine (Fig. 10).

**Fig 10.** Histopathology and viral component test in lung tissue at day 7 post challenged.

## Discussion

In this study we have investigated that RBD targeted binding Abs correlate very strongly with virus-neutralizing activity in natural infection and vaccination [24]. This study showed that an RBD immunogen offers a target for rational vaccine design both immunologically and from a manufacturability point of view [25]. Our study highlights induction of robust anti-SARS-CoV-2 Ab responses and protection from SARS-CoV-2 infection and pathogenesis specifically when using the RBD + Alhydrogel + CpG vaccine formulation.

Our study gave consistent results with previous reports of recombinant protein Spike protein using CpG [26], nAbs were observed after two vaccinations in our study. However, these titers were superior compared to Inactivated vaccine that was used in this study as a control vaccine. Although the presence of neutralizing and non-neutralizing epitopes contributing to RBD is smaller compared to inactivated vaccine, using the Alhydrogel and CpG adjuvant, boost the immune response while also showing protective efficacy in NHP challenge study. A preclinical study aimed at developing the subunit vaccine using various adjuvants has shown most effective in inducing antibodies neutralizing pseudovirus and wild-type virus by using CpG 1018 and aluminum hydroxide as adjuvant [27]. We report higher live SARS-CoV-2 nAbs in group 4 of animals after the second vaccination with a relatively lower increase in binding Abs. Additionally, a smaller drop observed in cross neutralizing activity against the Delta strain selectively in animals vaccinated with RBD + Alhydrogel + GpG suggests potential maturation of antibody responses, and no significant result of animals with low and high CpG. While a role for non-neutralizing Abs in protection with vaccine-induced immunity is unclear, treatment with monoclonal Abs in therapeutic settings highlights benefits with Fc effector functions in reducing lung pathology [28].

A protective SARS-CoV-2 vaccine should ideally induce a protective mucosal response. We observed strong anti-RBD Ab responses in nasal, BAL and rectal secretions when vaccinating via the IM route, which may contribute substantially to minimizing transmission and onset of disease followed by a reduction in fecal shedding of virus respectively. It is unknown whether an intranasal route of vaccination will further improve mucosal immunity with local IgA Abs. Most SARS-CoV-2 vaccines approved under emergency use authorization follow a two dose regiment. Since we observed protective efficacy after two vaccinations, additional investigation including testing in humans with two vaccine doses is critical to assess protective efficacy. Notably, robust induction of cross-CoV neutralizing Ab responses when vaccinating with a multimeric RBD displaying nanoparticle immunogen with the -alum and CpG has recently been reported [30]. These data are consistent with strong immunogenicity of multimeric nanoparticle protein-based construct used in the Novavax vaccine adjuvanted with Matrix-M [32]. Collectively, we conclude that RBD-Alum-CpG may offer significant promise in improving immunogenicity of both monomeric and multimeric RBD based SARS-CoV-2 vaccines.

Robust CD8+ T cell responses, nAbs and Th1 biased CD4+T cells could also support protective immunity against SARS-CoV-2 [33]. Specifically, where nAb responses can be sub-optimal against emerging variants of concern (VOCs) [20,21,32], conserved epitope-based CD8+ T cell responses could support protection. No evidence of induction of robust CD8+ T cell responses was observed when using the recently approved mRNA-1273 vaccine from Moderna, inactivated SARS-CoV-2 from Sinovac or a recombinant protein from Novavax in NHPs [32]. Of note, the saponin-based adjuvant used in the Novavax vaccine induces CD8+ T cell responses with protein immunogens in mice [34]. To date, SARS-CoV-2 specific CD8+ T cells have only been induced by viral vectored ChAd0X-1 and Ad.26/and or Ad5 in NHPs or with the BNT162b2, a mRNA vaccine in humans [19]. Induction of anti-RBD CD8+ T cell responses in our current study is in contrast to our inability to consistently induce anti COVID-19 cell responses in MCVCOV1901 when using TLR-9 agonist-based adjuvants [30]. It is possible that the RBD immunogen is enriched in epitopes for CD8+ T cells or preferentially targets cross-presenting DCs. However, as discussed above, RBD-containing whole S protein-based vaccines expressed in mammalian cells do not induce appreciable CD8+ T cell responses. Alternatively, unique glycosylation patterns of a yeast expressed RBD immunogen [35] that could improve targeting of lectin receptors on DC subsets [36] including monocyte-derived inflammatory DCs, may better induce CD8+ T cells. In summary, yeast expression of RBD in combination with a Th1 biasing Alhydrogel + CpG adjuvant likely led to targeting and differentiation of inflammatory DCs, supporting the induction of CD8+ cells that may improve vaccine efficacy against emerging SARS-CoV-2 variants.

Differentiation of classical to intermediate monocytes has been documented in Dengue virus infections in humans [37], as well as when using TLR-9 adjuvants in MVC-COV1901 [30]. Intermediate monocytes favor B cell differentiation to anti-body-secreting cells (ASCs) via secretion of IL-10 and B cell-activating factor (BAFF) [37]. SARS-CoV-2 is a single-stranded (ssRNA) virus with potential to target TLR-9 receptors. Our data supports that respiratory infection with SARS-CoV-2 leads to activation of blood monocytes. Precise immunological mechanisms by which either anti-RBD Ab or T cell responses induced by RBD+alum+CpG vaccine in respiratory mucosa contribute to blocking activation of peripheral blood monocytes are unclear. We hypothesize that anti-RBD Ab responses in nasal and BAL mucosa could be contributing to rapid clearance of virus via formation of immune complexes and perhaps signaling via Fc inhibitory receptors on innate immune cells as seen with mAbs in therapeutic settings [31]. Indeed, a larger role for monocytes/macrophages in BAL after SARS-CoV-2 challenge has been reported in both humans and MVC-COV1901[30].

The animal models have been used, including mice, Wistar rats, New Zealand rabbits, and Indonesian Macaca fascicularis. These animals have been widely used for toxicity and immunogenicity tests. On the other hand, mice are low cost, but unfortunately, mice, rats, and rabbits are not recommended for SARS-CoV-2 challenge testing, considering that the animals do not match some animal ACE2 amino acids with hACE2. Rabbits are susceptible to MERS-CoV infection but still do not show significant histopathological changes. In recent studies, however, higher SARS-CoV-2 virus doses have been used, and there is virus shedding in the nose, throat, and rectum, but infections with these high doses did not show clear clinical symptoms [38;39].

The use of NHP, Macaca fascicularis, as an animal model, is a Golden standard of animal model that is sensitive to SARS-CoV-2 infection and is also easily available in Indonesia. So far, Macaca fascicularis from Indonesia is widely available and used for research on drugs or other vaccines. SARS-CoV-2 infection in Macaca fascicularis will cause mild to moderate respiratory manifestations. This study with NHP was carried out at the Animal BSL-3 facilities belonging to the Professor Nidom Foundation. The Animal BSL-3 facilities have obtained a permit to operate as a COVID-19 diagnostic and research laboratory from the Ministry of Health of the Republic of Indonesia based on the Decree of the Minister of Health of the Republic of Indonesia in 2020 (SR.01.07/II/1859/2020 [40] and has been used for biohazard research on SARS-CoV-2 in stray cats and other pathogens, such as the H5N1 viruses; Nipah Virus [41;42;43] .

This study focuses on the acute phase of SARS-CoV-2 infection in Maccaca fascicularis. During the study, all animals infected with Wuhan-like and Delta viruses from Indonesian strain did not show clinical symptoms of the disease. This is following research by Urano et.al. [44] which showed all young Cynomolgus macaques appeared healthy without any clinical symptoms and research by Böszörményi et al [45] which only showed mild symptoms in macaques infected by SARS-CoV-2. Changes in hematological parameters and blood chemistry in Human COVID-19 are characterized by lymphopenia, thrombocytopenia, decreased hemoglobin, albumin, and increased ALT, AST, albumin, and creatinine in severe cases [46-47]. While changes in serum chemistry and hematological parameters can be important markers of disease, we found that SARS-CoV-2 infection in Maccaca fascicularis did not result in significant changes. These results are similar to studies in other NHPs on SARS-CoV-2 infection [44, 48-50]. It is difficult to conclude whether this change is related to SARS-CoV-2 infection because it was not followed by clinical changes or other parameters.

The RBD has been selected as an antigen candidate that can be produced at large scale with a thermostable (2-8 °C) formulation suitable for deployment in Low-middle countries. Our study supports the testing of an RBD-based immunogen adjuvanted with Alhydrogel and CpG in human trials that might offer a cost-effective, scalable and the most stable SARS-CoV-2 vaccine.

### Conclusions

Preclinical studies indicate that the recombinant COVID-19 vaccine has safe and shown immunogenic potential against the Indonesian SARS-COV-2 virus strain at a dose level of 12.5 µg and 25 µg/dose RBD adjuvanted with alhydrogel and CpG which corresponds to the non-human primate and human equivalent dose.

## Supporting information

Supplemental datas

## Acknowledgements

We want to express our heartfelt thanks to all the dedicated animal care personnel who made this research possible through their hard work. We are grateful to PT. Bio Farma for their financial support, which greatly assisted in our study. We would like to acknowledge Adriansjah Azhari as Head of Indonesian Health Institute of PT. Bio Farma for his vital role in project support, Maria Elena Bottazi and Peter Hotez from Baylor College of Medicine for providing the seed and support during development of COVID-19 vaccine, and Dynavax for providing the CpG. These contributions were crucial in our research, and we are thankful for the collaboration and support that made this study successful.

