## Supplemental datas for "Study of the immunogenicity, efficacy and safety of recombinant RBD SARS-CoV-2 vaccine with CpG adjuvant in rodent, non-rodent and Maccaca fascicularis using Indonesian Strain Virus"

A. SUPPLEMENTARY DATA OF ACUTE TOXICITY STUDY IN WISTAR RATS

Table SA.1. The clinical symptoms and local reactions of acute toxicity study in Wistar Rats

| Day- | Saline | Alum 750 µg | CpG 1500 µg | Alum 750µg + CpG 1500µg | RBD 25µg + Alum 750µg + CpG 750µg | RBD 25µg + Alum 750µg + CpG 1500µg | RBD 12,5µg + Alum 750µg + CpG 750µg | RBD 12,5µg + Alum 750µg + CpG 1500µg |
|---|---|---|---|---|---|---|---|---|
| 1 | Normal | Normal | Normal | Normal | Normal | Normal | Normal | Normal |
| 2 | Normal | Normal | Normal | Normal | Normal | Normal | Normal | Normal |
| 3 | Normal | Normal | Normal | Normal | Normal | Normal | Normal | Normal |
| 4 | Normal | Normal | Normal | Normal | Normal | Normal | Normal | Normal |
| 5 | Normal | Normal | Normal | Normal | Normal | Normal | Normal | Normal |
| 6 | Normal | Normal | Normal | Normal | Normal | Normal | Normal | Normal |
| 7 | Normal | Normal | Normal | Normal | Normal | Normal | Normal | Normal |
| 8 | Normal | Normal | Normal | Normal | Normal | Normal | Normal | Normal |
| 9 | Normal | Normal | Normal | Normal | Normal | Normal | Normal | Normal |
| 10 | Normal | Normal | Normal | Normal | Normal | Normal | Normal | Normal |
| 11 | Normal | Normal | Normal | Normal | Normal | Normal | Normal | Normal |
| 12 | Normal | Normal | Normal | Normal | Normal | Normal | Normal | Normal |
| 13 | Normal | Normal | Normal | Normal | Normal | Normal | Normal | Normal |
| 14 | Normal | Normal | Normal | Normal | Normal | Normal | Normal | Normal |

Clinical symptoms are observed by observing the animal's behaviour, the local reactions are observed by observing the injection site.

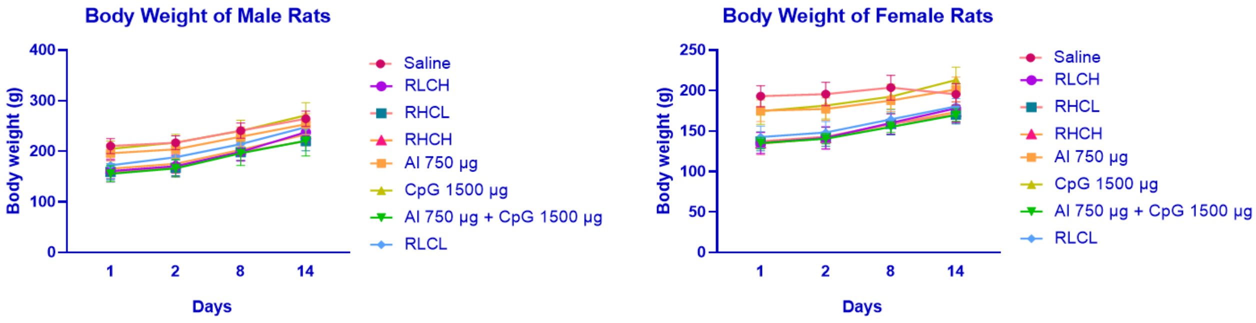


Figure S A.1.Profile of body weight changes of male and female rats after vaccination in acute toxicity study. (A) Female rats (n=10); (B) Male rats (n=10). Animals were observed for body weight changes after vaccination (days 1, 2, 8 and 14).

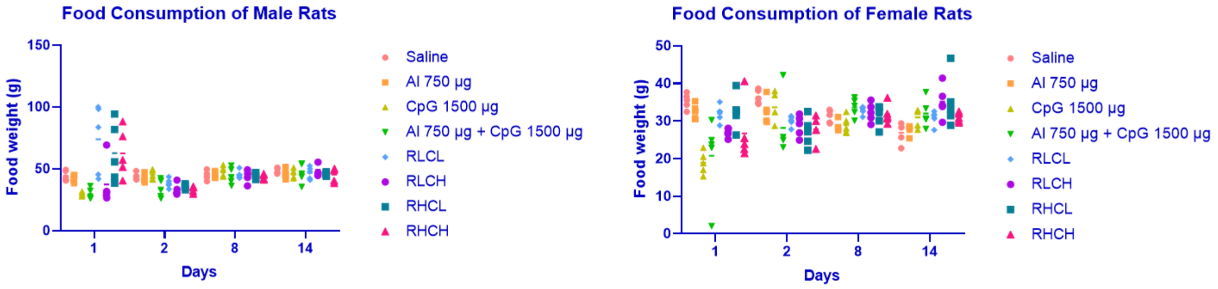


Figure S A.2.Profile of food consumption rats in acute toxicity study. Male rats (n=10) and Female rats (n=10). Animals were observed on day 1, 2, 8, and 14. Control represents animals that received a saline vaccination.

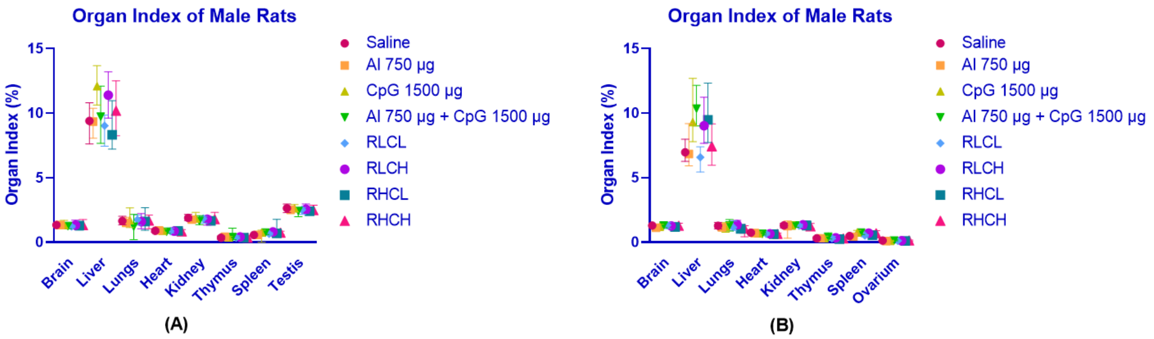


Figure S.A.3 Profile of rats' organ index in acute toxicity study. (A) Male rats (n=10); (B) Female rats (n=10).

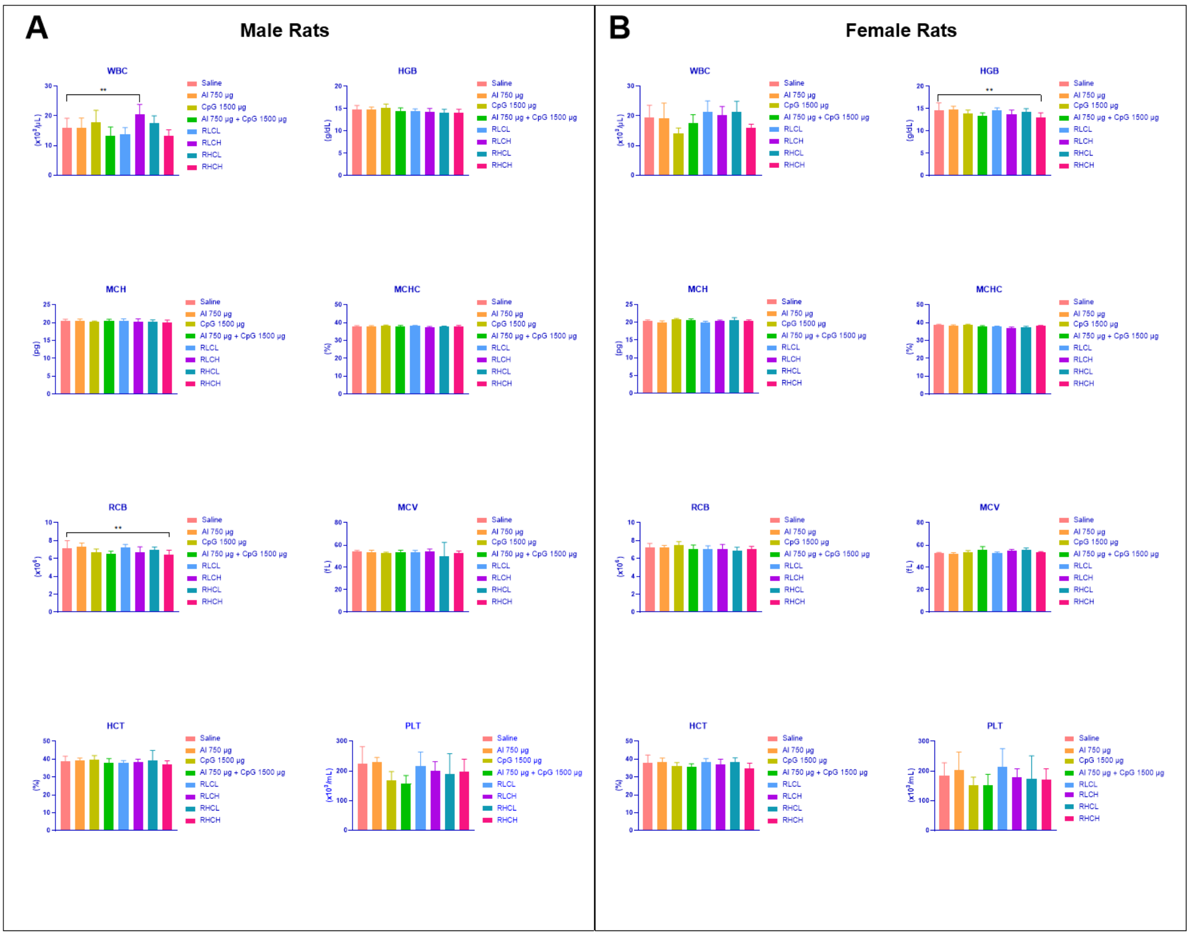


Figure S A.4. Hematological analysis of vaccinated rats in acute toxicity study.

Control represents animals that received a saline vaccination. There were no statistically significant differences (ns>0.05) for haematological analysis of MCH, MCHC, MCV, HCT and PLT on day-14th after vaccination compared to control. The significant differences were determined using ordinary one-way ANOVA followed by Dunnett multiple comparison test (ns>0.05; *p<0.05; **p<0.01; ***p<0.001; ****p<0.0001). (A) Hematological analysis on male rats; (B) Hematological analysis on female rats

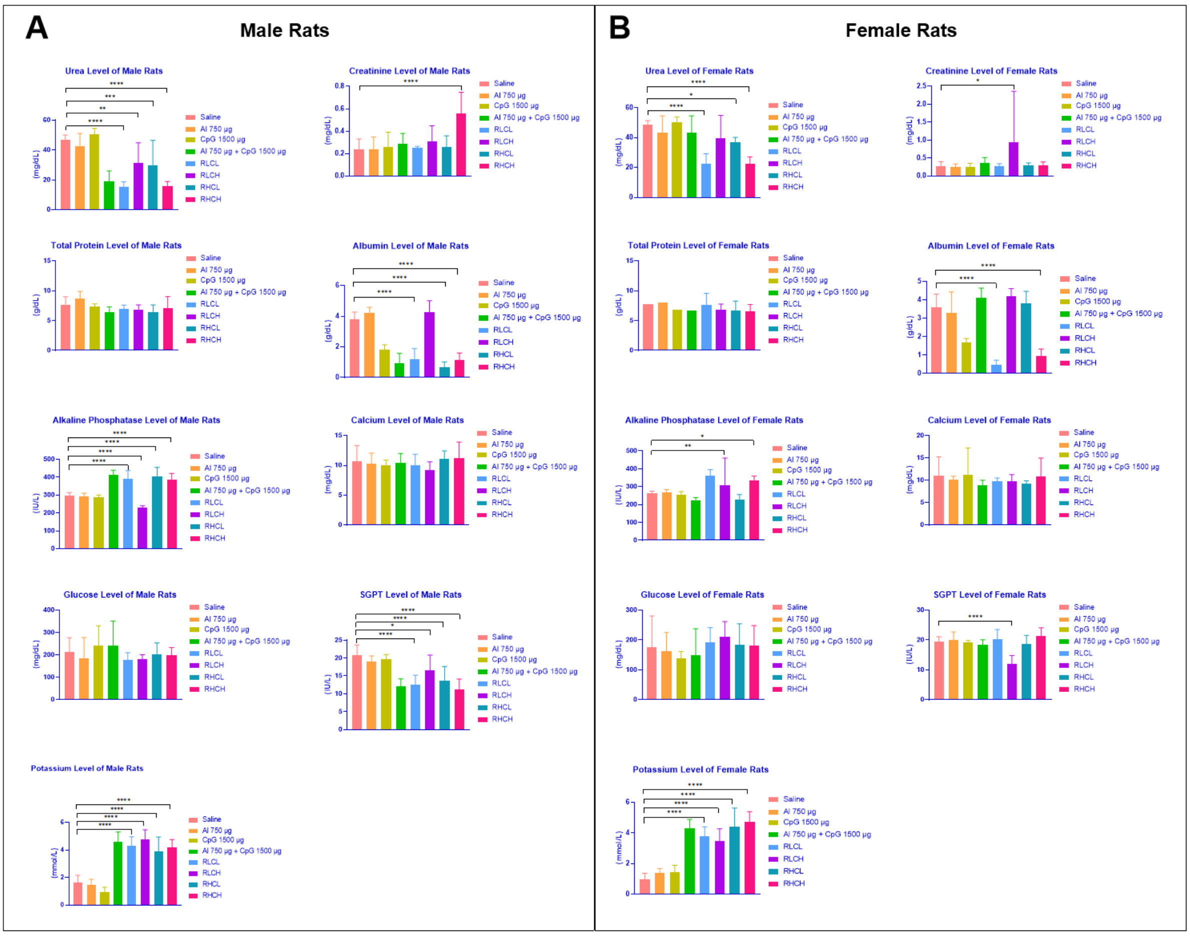


Figure S.A.5 Blood chemistry analysis of vaccinated rats in acute toxicity study.
Control represents animals that received a saline vaccination. Blood samples were collected on day 14. The significant differences were determined using one-way ANOVA followed by Dunnett multiple comparison test (ns>0.05; *p<0.05; **p<0.01; ***p<0.001; ****p<0.0001). (A) Blood chemistry analysis on male rats(n=10); (B) Blood chemistry analysis on female rats (n=10)

### B. SUPPLEMENTARY DATA OF SUBCHRONIC TOXICITY STUDY IN WISTAR RATS

Table S.B.1 The clinical symptoms and local reactions of subchronic toxicity study in Wistar Rats

| **Day-** | **Saline** | **Alum 750 µg** | **CpG 1500 µg** | **Alum 750µg + CpG 1500µg** | **RBD 25µg + Alum 750µg + CpG 750µg** | **RBD 25µg + Alum 750µg + CpG 1500µg** | **RBD 12,5µg + Alum 750µg + CpG 750µg** | **RBD 12,5µg + Alum 750µg + CpG 1500µg** |
|---|---|---|---|---|---|---|---|---|
| 1 | Normal | Normal | Normal | Normal | Normal | Normal | Normal | Normal |
| 2 | Normal | Normal | Normal | Normal | Normal | Normal | Normal | Normal |
| 3 | Normal | Normal | Normal | Normal | Normal | Normal | Normal | Normal |
| 4 | Normal | Normal | Normal | Normal | Normal | Normal | Normal | Normal |
| 5 | Normal | Normal | Normal | Normal | Normal | Normal | Normal | Normal |
| 6 | Normal | Normal | Normal | Normal | Normal | Normal | Normal | Normal |
| 7 | Normal | Normal | Normal | Normal | Normal | Normal | Normal | Normal |
| 8 | Normal | Normal | Normal | Normal | Normal | Normal | Normal | Normal |

| | | | | | | | | |
|---|---|---|---|---|---|---|---|---|
| 14 | Normal | Normal | Normal | Normal | Normal | Normal | Normal | Normal |
| 21 | Normal | Normal | Normal | Normal | Normal | Normal | Normal | Normal |
| 28 | Normal | Normal | Normal | Normal | Normal | Normal | Normal | Normal |
| 35 | Normal | Normal | Normal | Normal | Normal | Normal | Normal | Normal |
| 42 | Normal | Normal | Normal | Normal | Normal | Normal | Normal | Normal |
| 49 | Normal | Normal | Normal | Normal | Normal | Normal | Normal | Normal |
| 56 | Normal | Normal | Normal | Normal | Normal | Normal | Normal | Normal |
| 59 | Normal | Normal | Normal | Normal | Normal | Normal | Normal | Normal |

Clinical symptoms are observed by observing the animal's behaviour, the local reactions are observed by observing the injection site.

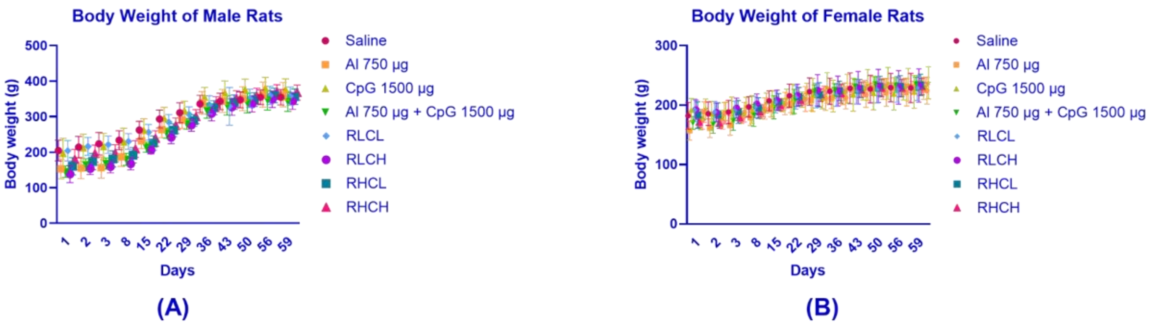


Figure S.B.1.Profile of body weight changes of male and female rats in subchronic toxicity study. (A) Male rats (n=10); (B) Female rats (n=10). Animals were observed for body weight changes on day 1, 2, 3, 8, 15, 22, 29, 36, 43, 50, 56, and 59.

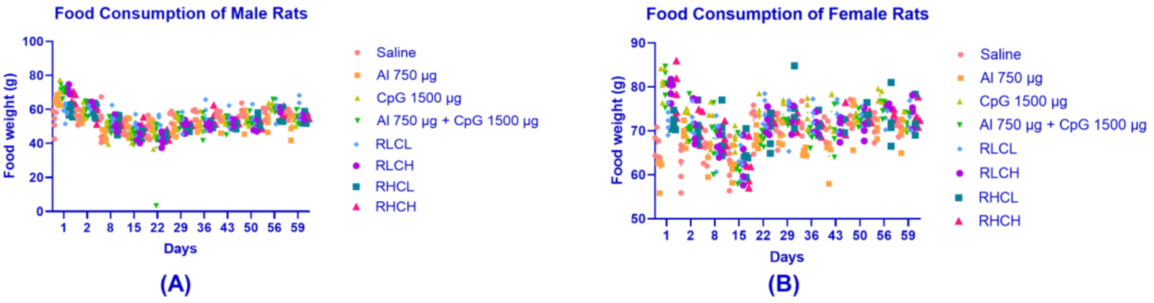


Figure S.B.2 Profile of rats' food consumption subchronic toxicity study. (A) Male rats (n=10); (B) Female rats (n=10). Animals were observed on day 1, 2, 8, 15, 22, 29, 36, 43, 50, 56, and 59.

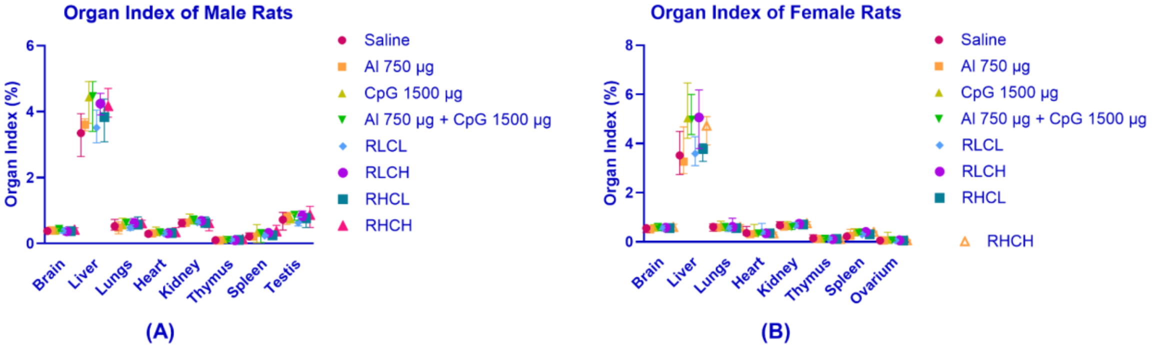


Figure S.B.3 Profile of rats' organ index in subchronic toxicity study. (A) Male rats (n=10); (B) Female rats (n=10).

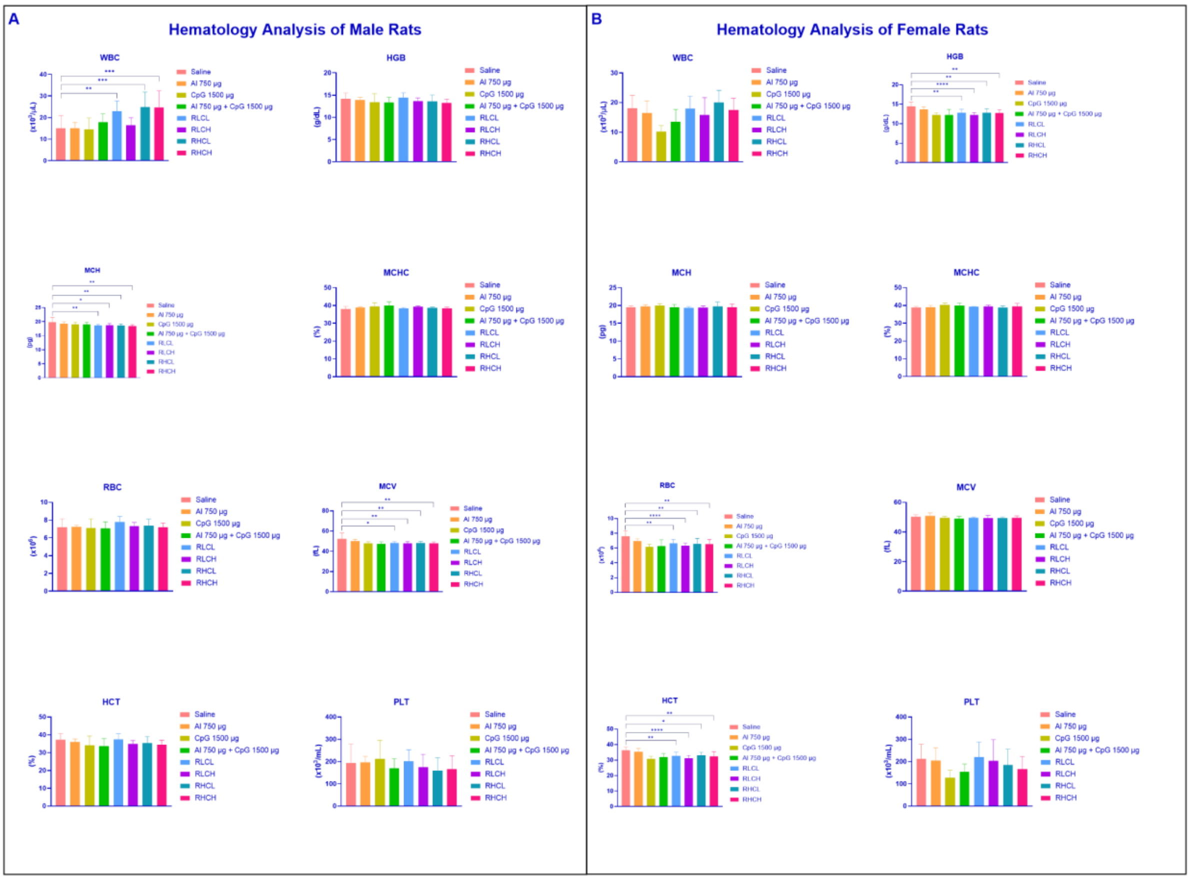


Figure S B.4. Hematological analysis of vaccinated rats in sub chronic toxicity study.

Control represents animals that received a saline vaccination. Blood samples were collected on day 59. There were no statistically significant differences (ns>0.05) for haematological analysis of HGB, MCHC, RBC, HCT, and PLT in male rats. On the other hand, there were no statistically significant differences of WBC, MCH, MCHC, MCV, and PLT in female rats. The significant difference was determined using one-way ANOVA followed by Dunnett multiple comparison test (ns>0.05; *p<0.05; **p<0.01; ***p<0.001; ****p<0.0001). (A) Hematological analysis on male rats; (B) Hematological analysis on female rats

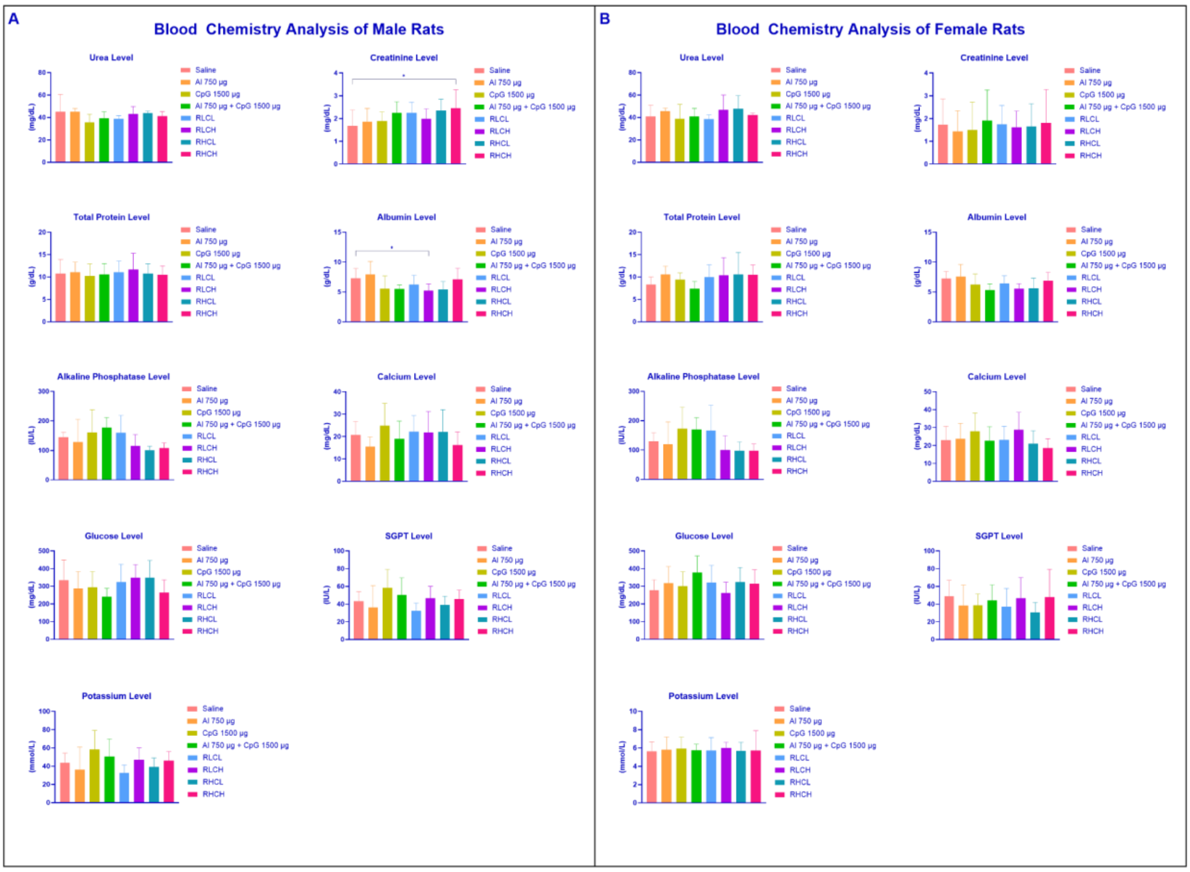


Figure S.B.5 Blood chemistry analysis of vaccinated rats in sub chronic toxicity study.

Control represents animals that received a saline vaccination. Blood samples were collected on day 59. There were no statistically significant differences (ns>0.05) for blood chemistry analysis of urea, total protein, albumin, alkaline phosphatase, calcium, glucose, SGPT, and potassium level in male rats compared to control. While there were no statistically significant differences were determined in all parameters of female rabbits The significant differences was determined using one-way ANOVA followed by Dunnett multiple comparison test (ns>0.05; *p<0.05; **p<0.01; ***p<0.001; ****p<0.0001). (A) Blood chemistry analysis on male rats(n=10); (B) Blood chemistry analysis on female rats (n=10)

C. SUPPLEMENTARY DATA OF ACUTE TOXICITY STUDY IN NEW ZEALAND RABBITS

Table S.C.1. The clinical symptoms and local reactions of acute toxicity study in New Zealand Rabbit

| Day- | Saline | Alum 750 µg | CpG 1500 µg | Alum 750µg + CpG 1500µg | RBD 25µg + Alum 750µg + CpG 750µg | RBD 25µg + Alum 750µg + CpG 1500µg | RBD 12,5µg + Alum 750µg + CpG 750µg | RBD 12,5µg + Alum 750µg + CpG 1500µg |
|---|---|---|---|---|---|---|---|---|
| 1 | Normal | Normal | Normal | Normal | Normal | Normal | Normal | Normal |
| 2 | Normal | Normal | Normal | Normal | Normal | Normal | Normal | Normal |
| 3 | Normal | Normal | Normal | Normal | Normal | Normal | Normal | Normal |
| 4 | Normal | Normal | Normal | Normal | Normal | Normal | Normal | Normal |
| 5 | Normal | Normal | Normal | Normal | Normal | Normal | Normal | Normal |
| 6 | Normal | Normal | Normal | Normal | Normal | Normal | Normal | Normal |
| 7 | Normal | Normal | Normal | Normal | Normal | Normal | Normal | Normal |
| 8 | Normal | Normal | Normal | Normal | Normal | Normal | Normal | Normal |
| 9 | Normal | Normal | Normal | Normal | Normal | Normal | Normal | Normal |
| 10 | Normal | Normal | Normal | Normal | Normal | Normal | Normal | Normal |
| 11 | Normal | Normal | Normal | Normal | Normal | Normal | Normal | Normal |
| 12 | Normal | Normal | Normal | Normal | Normal | Normal | Normal | Normal |
| 13 | Normal | Normal | Normal | Normal | Normal | Normal | Normal | Normal |
| 14 | Normal | Normal | Normal | Normal | Normal | Normal | Normal | Normal |

Clinical symptoms are observed by observing the animal's behaviour, the local reactions are observed by observing the injection site.

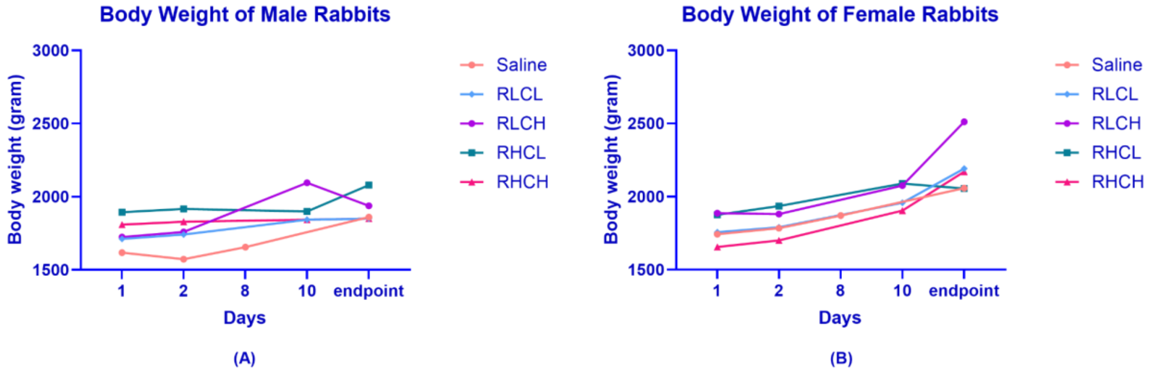


Figure S.C.1.Profile of body weight changes of male and female rabbits in acute toxicity study. (A) Male rabbits (n=5); (B) Female rabbits (n=5).

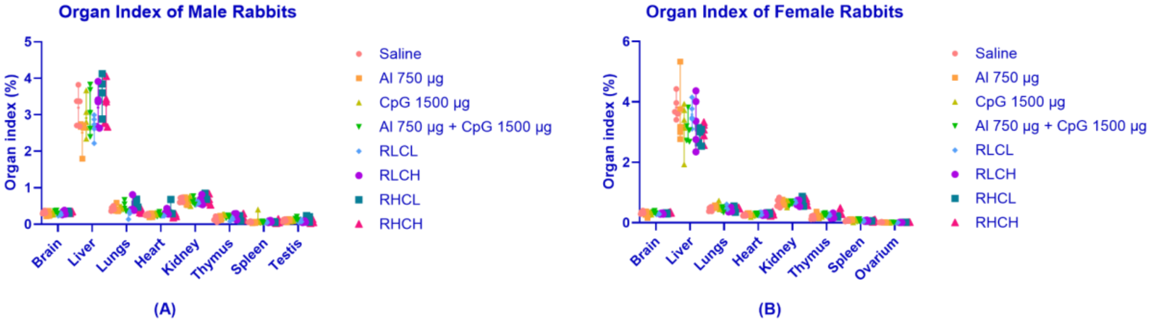


Figure S.C.2 Profile of rabbits' organ index in acute toxicity study. (A) Male rabbits (n=5); (B) Female rats (n=5).

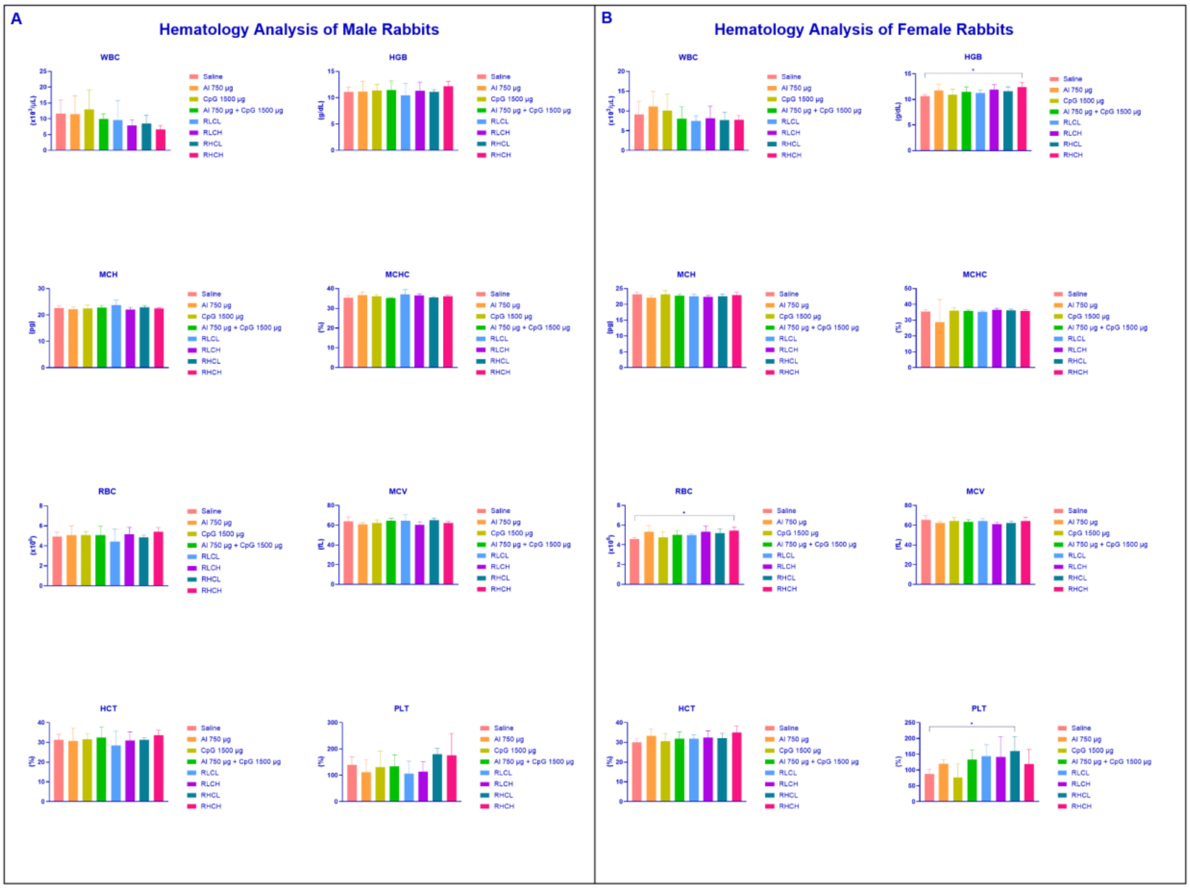


Figure S.C.3. Hematological analysis of vaccinated rabbits in acute toxicity study. There were no statistically significant differences (ns>0.05) for haematological parameters of male rabbits. While there were statistically differences in HGB and RBC of the RHCH group, and PLT of RHCL group. The significant differences were determined using one-way ANOVA followed by Dunnett multiple comparison test (ns>0.05; *p<0.05; **p<0.01; ***p<0.001; ****p<0.0001). (A) Blood chemistry analysis on male rabbits (n=5); (B) Blood chemistry analysis on female rabbits (n=5)

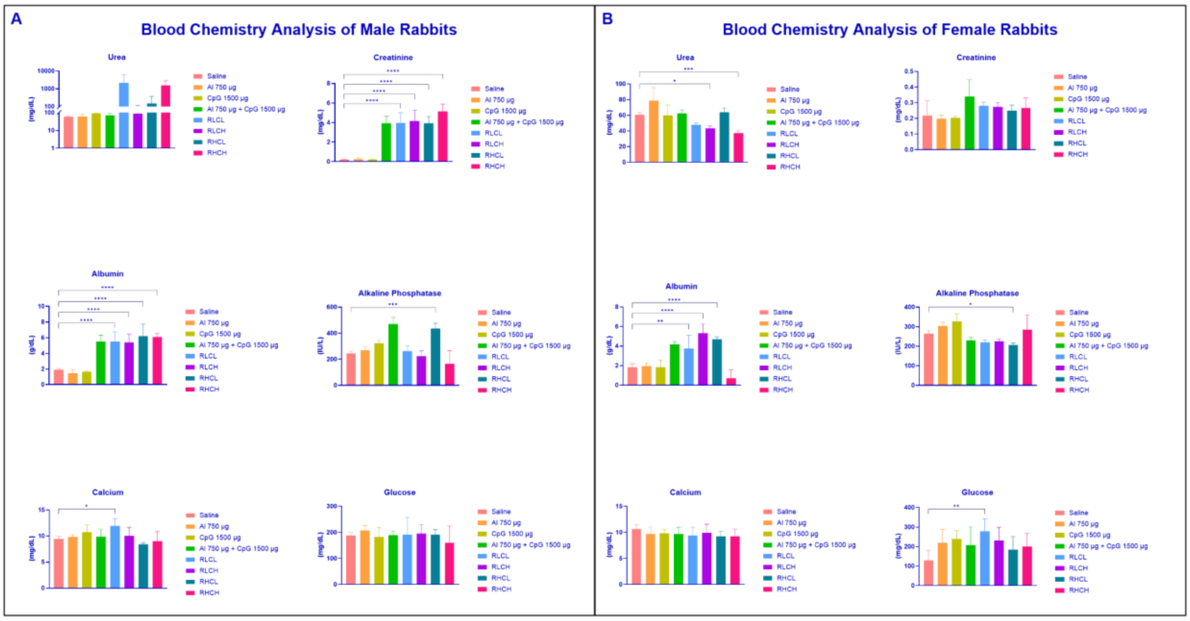


Figure S.C.4. Blood chemistry analysis of vaccinated rabbits in acute toxicity study. Control represents animals that received a saline vaccination. There were several statistically significant differences *(ns>0.05)* for blood chemistry analysis of urea, creatinine, albumin, alkaline phosphatase, calcium, and glucose concentration. The significant differences were determined using one-way ANOVA (*ns>0.9999; *p<0.05; **p<0.01; ***p<0.001; ****p<0.0001*). (A) Blood chemistry analysis on male rabbits; (B) Blood chemistry analysis on female rabbits

D. SUPPLEMENTARY DATA OF SUBCHRONIC TOXICITY STUDY IN NEW ZEALAND RABBITS

Table S.D.1 The clinical symptoms and local reactions of subchronic toxicity study in New Zealand rabbits

| Day- | Saline | Alum 750 µg | CpG 1500 µg | Alum 750µg + CpG 1500µg | RBD 25µg + Alum 750µg + CpG 750µg | RBD 25µg + Alum 750µg + CpG 1500µg | RBD 12,5µg + Alum 750µg + CpG 750µg | RBD 12,5µg + Alum 750µg + CpG 1500µg |
|---|---|---|---|---|---|---|---|---|
| 1 | Normal | Normal | Normal | Normal | Normal | Normal | Normal | Normal |
| 2 | Normal | Normal | Normal | Normal | Normal | Normal | Normal | Normal |
| 3 | Normal | Normal | Normal | Normal | Normal | Normal | Normal | Normal |
| 4 | Normal | Normal | Normal | Normal | Normal | Normal | Normal | Normal |
| 5 | Normal | Normal | Normal | Normal | Normal | Normal | Normal | Normal |
| 6 | Normal | Normal | Normal | Normal | Normal | Normal | Normal | Normal |
| 7 | Normal | Normal | Normal | Normal | Normal | Normal | Normal | Normal |
| 8 | Normal | Normal | Normal | Normal | Normal | Normal | Normal | Normal |
| 14 | Normal | Normal | Normal | Normal | Normal | Normal | Normal | Normal |
| 21 | Normal | Normal | Normal | Normal | Normal | Normal | Normal | Normal |
| 28 | Normal | Normal | Normal | Normal | Normal | Normal | Normal | Normal |
| 35 | Normal | Normal | Normal | Normal | Normal | Normal | Normal | Normal |
| 42 | Normal | Normal | Normal | Normal | Normal | Normal | Normal | Normal |
| 49 | Normal | Normal | Normal | Normal | Normal | Normal | Normal | Normal |
| 56 | Normal | Normal | Normal | Normal | Normal | Normal | Normal | Normal |
| 59 | Normal | Normal | Normal | Normal | Normal | Normal | Normal | Normal |

Clinical symptoms are observed by observing the animal's behaviour, the local reactions are observed by observing the injection site.

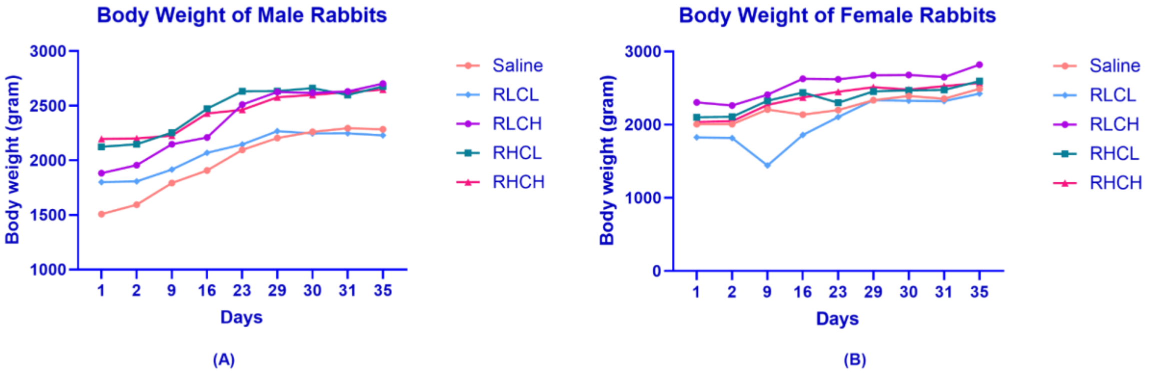


Figure S.D.1.Profile of body weight changes of sub chronic toxicity study in New Zealand rabbits. (A) Male rabbit; (B) Female rabbits. Animals were observed for body weight changes on day 1, 2, 9, 16, 23, 29, 30, 31, and 35.

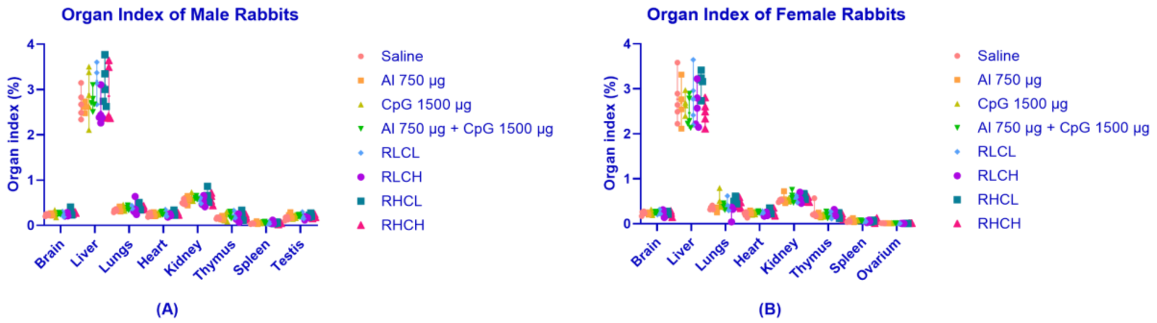


Figure S.D.2 Profile of rabbits' organ index in sub chronic toxicity study. Male rabbits (n=5); (B) Female rabbits (n=5)

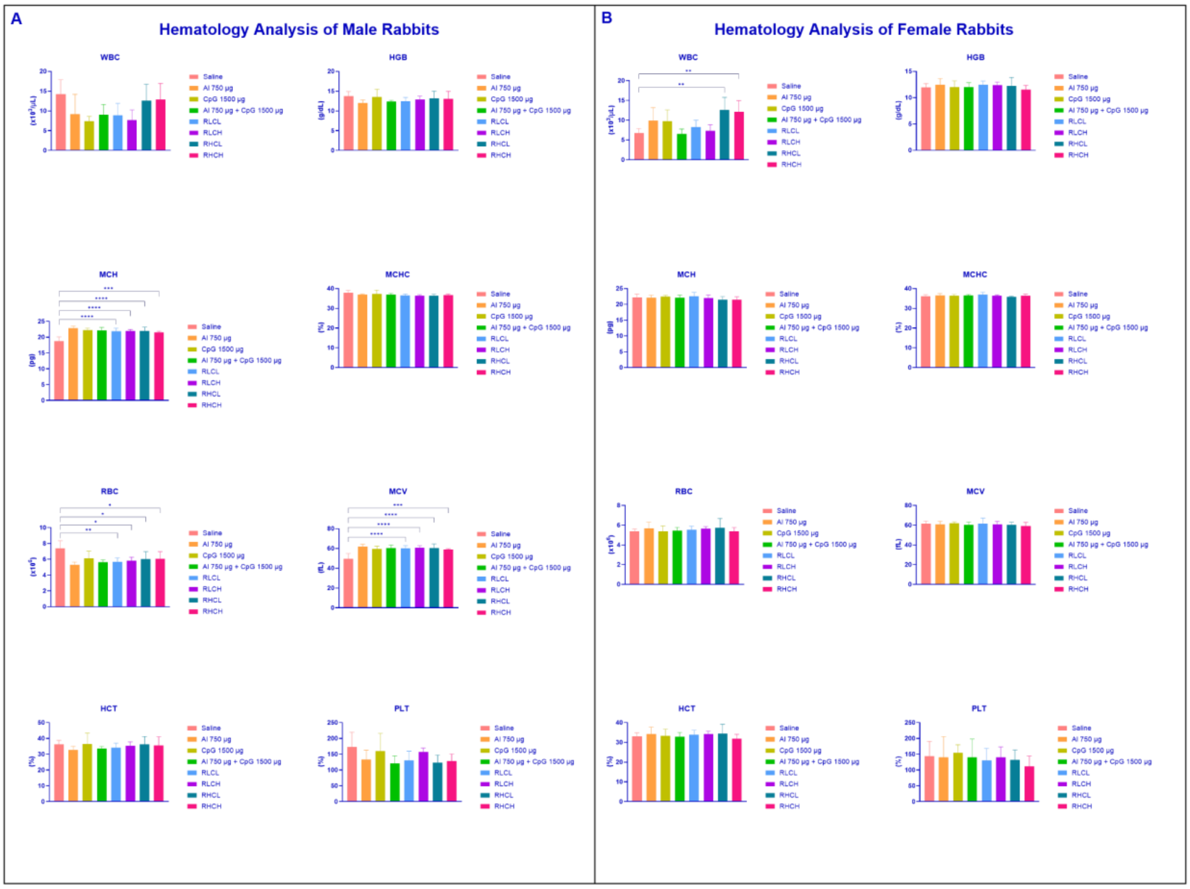


Figure S.D.3. Hematological analysis of vaccinated rabbits in sub chronic toxicity study.

Control represents animals that received a saline vaccination. There were no statistically significant differences (ns>0.05) for haematological analysis of WBC, HGB, MCHC, PCT, and PLT in male rabbits. On the other hand, there were no statistically significant differences of HGB, MCH, MCHC, RBC, MCV, HCT, and PLT in female rabbits. The significant differences were determined using one-way ANOVA followed by Dunnett multiple comparison test (ns>0.05; *p<0.05; **p<0.01; ***p<0.001; ****p<0.0001). (A) Hematological analysis on male rabbits; (B) Hematological analysis on female rabbits

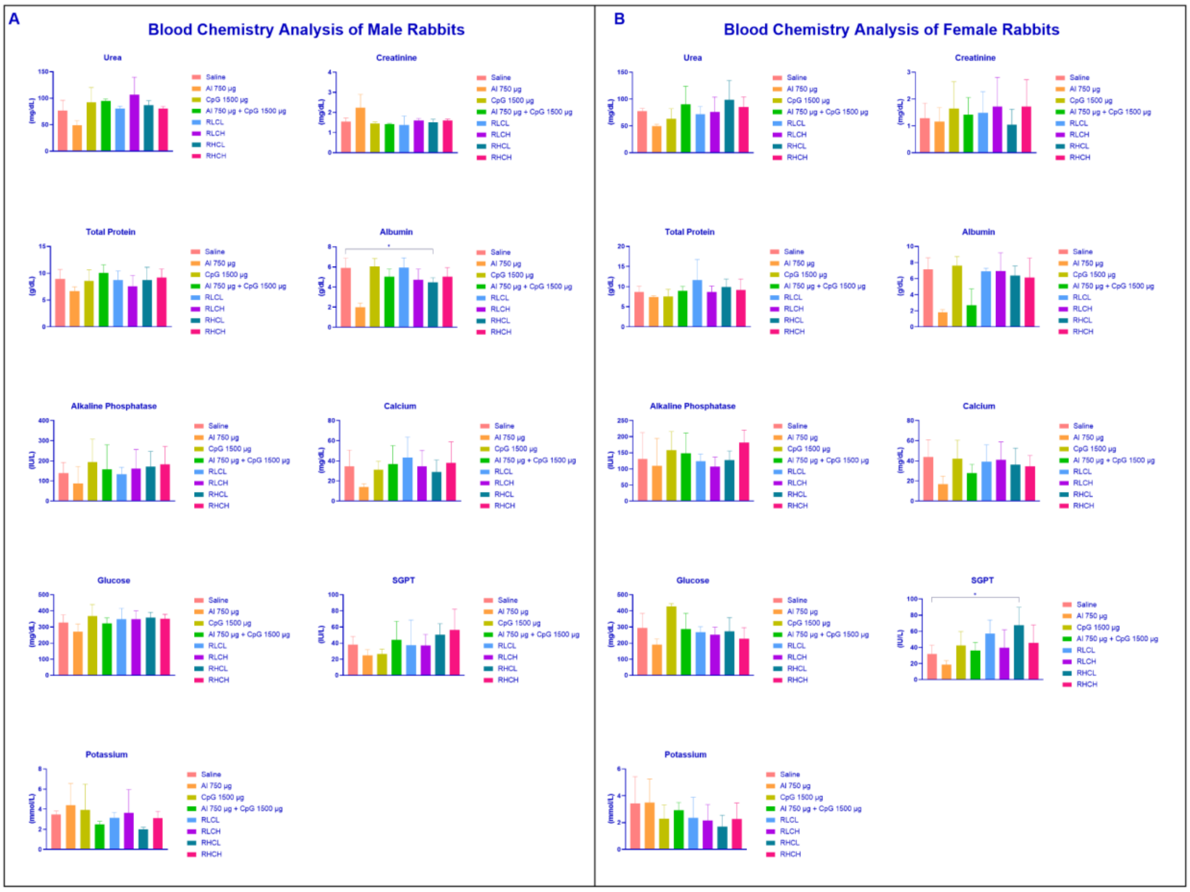


Figure S.D.4 Blood Chemistry analysis of vaccinated rabbits in sub chronic toxicity study.

Control represents animals that received a saline vaccination. There were no statistically significant differences (ns>0.05) for blood chemistry analysis of urea, creatinine, total protein, alkaline phosphatase, calcium, glucose, SGPT, and potassium concentration in male rabbits compared to control. While there were no statistically significant differences were determined in urea, creatinine, total protein, albumin, alkaline phosphatase, calcium, glucose, and potassium concentration female rabbits. The significant differences was determined using one-way ANOVA followed by Dunnett multiple comparison test (ns>0.05; *p<0.05; **p<0.01; ***p<0.001; ****p<0.0001). (A) Blood chemistry analysis on male rabbits; (B) Blood chemistry analysis on female rabbits
